# Modulation of H3K4 trimethylation by KDM5A and MLLs impacts metabolic adaptability in prostate and cervical cancer cells

**DOI:** 10.1101/2024.05.02.592178

**Authors:** R. Kirtana, Soumen Manna, Samir Kumar Patra

## Abstract

Chemical modifications of chromatin modulate gene expression and induce essential metabolic plasticity for tumor growth. Accumulation of H3K4me3 in the promoter of a gene activates transcription by making the promoter accessible to the polymerases. Methylation of H3K4 is catalysed by MLLs and demethylation of H3K4me3 is catalysed by KDM5 family proteins. Herein, we investigated if genes encoding the enzymes involved in glucose metabolism are dependent on KDM5A and MLL1, and if targeting the H3K4me3 would help in modulating the resilience of cancer cells. We present that KDM5A modulates most of the metabolic genes in a demethylase dependent manner as assesses by H3K4me3 occupancy on G6PD and catalase promoters. Targeting its expression would indeed help in sensitizing cancer cells to ROS dependent apoptotic cell death. We elucidated the differences in the epigenetic regulation in cancerous cells originated from cervical and prostate tissues and used a normal skin keratinocyte for comparison. In cervical and prostate cancers - KDM5A activated glycolysis but downregulates other metabolic processes. In cervical cancer, which majorly depends on PPP, changes in KDM5A did not modulate the G6PD expression. Further, we have shown that curcumin treatment enhanced KDM5A expression and downregulated MLL2 in cancer cell lines but not in keratinocyte cells. Curcumin inhibited metabolic pathways and enhanced apoptosis in cancer cells without affecting keratinocyte cells by modulating KDM5A and MLL levels. This work also strengthens the basic concept that, epigenetic modulations of genes in a tissue precisely depends on signal and sites of modification(s).

## Introduction

Mitochondria regulates vital networking by harboring many catabolic and anabolic reactions and contribute to retrograde signaling (by generating different reactive oxygen species (ROS); and indirectly regulate gene expression by providing co-factors like acetyl-CoA and α-ketoglutarate (α-KG) to epigenetic modifiers thereby regulating their function; and mediate apoptotic pathways as well [1–6]. Generally, cancer cells derail from OXPHOS dependence to glycolysis and pentose phosphate pathway (PPP). The tumor microenvironment is a major factor in dictating the metabolic adaptability of tumor cells. With this phenotypic plasticity on metabolic regulation, cancer cells attain oxidative and chemotherapeutic resistance [7–8].

The control over gene expression is majorly governed by transcription factors and epigenetic modulators. Epigenetic regulation of mitochondrial protein synthesis is crucial factor during cancer initiation and progression, thus targeting the epigenetic factors is a promising approach to either restore or inhibit mitochondrial activity depending on the metabolic pathway controlling the tumor growth and the stage of cancer(s) [9]. Although many summaries acknowledge how metabolic processes could modify the chromatin by donating co-factors to the writers, chromatin remodeler and erasers [10–13] and some suggested targeting metabolic pathways to target cancer [14], not much is explored on how epigenetic players regulate the genes involved in the metabolic processes. Some of the epigenetic enzymes have been reported to regulate genes involved in metabolism, like LSD1 repressing gluconeogenic genes; DNMT1 silences FBP1and KDM3A activates GLUT1 [15–17]. Many aspects need to be deciphered on how epigenetic mediators can control metabolic regulatory genes while attaining plasticity during cancer progression.

KDM5 family enzymes are histone demethylases that act on the transcriptional active mark H3K4me3 [18]. Roles of KDM5A in controlling cellular metabolic processes are emerging. Most of the metabolism related gene promoters harbor this active mark for constitutive expression of the downstream gene. Downregulation of KDM5A was observed during differentiation of MEFs into myocyte with increased respiratory rate and mitochondrial function, proving that KDM5A regulates these genes in a demethylase dependent manner [19]. Roesch et al., [7] and Sharma et al., [20] identified a subset of slow growing metabolically distinct population of cells that overexpress KDM5A/B and derive ATP by OXPHOS and not aerobic glycolysis and increased mitochondrial activity was associated with enhanced H2O2 levels [7]. In melanoma cells with KDM5B overexpression, an increase in glucose 6 phosphate dehydrogenase (G6PD) mediated shift from glycolytic phenotype to PPP (for antioxidant NADPH) was observed [21]. Secombe group [**22, 23** and references therein], have depicted that Drosophila KDM5 is a critical regulator of mitochondrial function in a demethylase independent manner by using PHD3 domain that recognize H3K4me2/3 and activate gene expression, and they concluded that in adult flies, KDM5 is primarily an activator of gene expression. Comparing mitochondria from myofibrils of wild type and hypomorphic mutants showed morphological defects with decreased ATP levels. The mutant flies also showed increased superoxide levels, high citrate levels and enhanced expression of TCA cycle enzymes in an indirect manner. Lopez-Bigas et al., (2008) [24] showed that in SAOS-2 cells lacking pRB (a KDM5A interacting protein), KDM5A knockdown led to long rod-shaped mitochondria. In contrast to the observations in Drosophila mutants, these knockdown cells did not exhibit any changes in superoxide levels. Also, in mammalian cells (Lopez-Bigas et al [24]), mitofusion-2 (MFN2) was noted to be a direct target of KDM5A, but the Drosophila paralog (Marf) was unregulated leading to a hypothesis that this regulation is probably tissue-specific [22]. In Drosophila, KDM5 was defined as an activator of genes involved in maintenance of redox homeostasis by interacting with Foxo (the stress resistance transcriptional factor) independent of its enzymatic activity by recruiting HDAC4 for deacetylation of Foxo, and absence of KDM5 attenuates acetylated Foxo recruitment to its target gene promoters. Absence of KDM5C enhanced oxidative stress mediated damage of proteins and DNA leading to cell death [23].

In view of this and additional literature survey, it was hypothesized that KDM5A can either be an activator or repressor of metabolic genes depending on the tissue type and developmental stage, which might further be governed by its interaction with other cellular proteins, subject to their availability. Secombe’s laboratory proved that male and female flies react differently towards KDM5 mutation (demethylase dead). Male flies expressing the KDM5 mutant were sensitive to paraquat (inducer of oxidative stress) when compared to females, and their gene expression also varied. Thus, we sought to understand the sex-specific role of KDM5A using a female originated cervical cell line (HeLa) and a male originated prostate cancer cell line (PC3). HaCaT (the normal cell line without mutational load) was used to compare the changes following the treatments.

Through this study, by modulating the expression of a writer (MLL1) and an eraser (KDM5A) enzyme, we tested the hypothesis whether H3K4me3 controls the metabolic fate in cervical and prostate cancer cells. We further examined whether targeting H3K4me3 demethylation may provide a novel way of reprogramming the cancer cell metabolic adaptability by utilizing some edible phytochemical compounds that act through regulating the expression of epigenetic modifiers thereby inhibiting the tumor growth by modifying their nature of adaptability and metabolic plasticity.

## Materials and Methods

### Gene expression profiling and correlation analysis

The UALCAN database was used to assess the KDM5A transcript profile in cervical and prostate tumor tissues relative to its expression in normal tissue. The expression profile of KDM5A across multiple tumor samples paired with the normal tissue was analyzed using gepia2. Using panther GO search for key metabolic pathways like glycolysis, TCA cycle, ATP synthesis and PPP along with genes involved in ROS scavenging and lactate fermentation (listed from literature review), we retrieved 80 metabolic signature genes for comparison with KDM5A expression in cervical squamous cell carcinoma (CESC) and prostate adenocarcinoma (PRAD) datasets. Further to study if KDM5A, MLLs and H3K4me3 occupy the promoters under our study, ChIP-seq data (from breast cancer cell lines) was retrieved from ChIP-atlas and analyzed using IGV [21, 25].

### Cell Culture

HeLa and HaCaT cells were cultured in MEM and DMEM media respectively, with 10% FBS (Gibco-10270106) and 1% anti-anti (Gibco-15240096). PC3 was grown in F12 media supplemented with 10% FBS, 1% anti-anti and L-glutamine (Gibco -15240-062). During treatments, depending on cell doubling size, seeding density was calculated to attain 70% confluency within 24hrs, following transfections or drug incubations. For microscopy experiments, seeding density was reduced to avoid overcrowding.

### siRNA transfection

In HeLa and HaCaT cell lines, both knockdown and overexpression was performed for 48hrs, but in PC3 the treatment duration was 24hrs. Transfection was attained using lipofectamine 3000 (Invitrogen L3000-15) following manufacturer’s instructions. Plasmid concentration varied from 6 well to 60mm plate and we used 5 or 10ug respectively to induce KDM5A overexpression. The pcDNA-triple epitope SFB-tagged-RBP2 (KDM5A) overexpression construct was borrowed from Dr. Shweta Tyagi (CDFD) [26].

Efficiency of knockdown and overexpression was confirmed by immunoblotting. As MLL1 is a huge protein (432KDa), and we could not perform immunoblotting successfully, so we analysed its knockdown efficiency by immunocytochemistry.

### MTT assay

Cell viability changes following drug treatment were determined by measuring absorbance off MTT (Himedia-TC191) in living cells. In brief, 24hrs prior to the drug administration, 10^3^ to 1.5*10^3^ cells were seeded into a 96-well plate, followed by replacing the normal media with drug dissolved media at required dilutions. The plate was then replaced into the incubator for 24hrs for the drug to be effective, followed by addition of MTT media, which was incubated for another 6hrs followed by DMSO (Himedia – AS121) mediated dissolution of the formazan crystals. A colorimetric analysis of the optical density at 570nm gave us the percentage of viable cells following treatment with different concentrations of a particular drug.

The percentage of viability was calculated as follows:

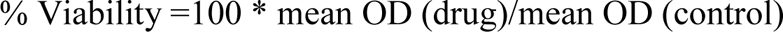

Drugs used in this study are listed in supplementary information. Cells were treated with these drugs for 24hrs prior to downstream experimentation.

### qRT-PCR experiments

To quantify the changes in gene expression profile following treatments, RT-PCR was performed. RNA isolation was performed by TRIzol (Thermo - 15596018) method, using isopropanol (himedia-MB063) for RNA precipitation and 70% alcohol (Himedia-MB228) to wash off excess salts and contaminants, and the RNA was dissolved in DEPC water. Reverse transcriptase reaction was setup to synthesize cDNA as per manufacturer’s instructions (Genesure – PGK163A) with the above isolated RNA with incubation at 42°C for 60min and 70°C for 5 min using a 20uL reaction mixture. Using this cDNA (1 ug) as template, and gene specific primers (300nM) respective to different physiological processes, RT-PCR was performed using SYBR-green technology (Thermo – A25742) with the following cycling conditions 50.0°C for 1:00; 96.0°C for 6:00 and [96.0°C for 0:10; 55.0°C for 0:30; 72.0°C for 1:00] for 40 cycles. The data was analysed according to Livak’s method (ddCT calculation) using either GAPDH or B-actin as control [27]. Primers used in the study are listed in supplementary data.

### Western blotting

Whole cell lysate prepared by RIPA lysis buffer (sigma –R0278-50ML) were electrophoresed on 8-12% SDS-PAGE gels depending on target proteins analysed as per previous protocol [28], resolved and transferred to nitrocellulose membrane (Axiva – 160300RI), blocked with 5% skim milk for an hour. All primary antibodies were prepared in 1% BSA (Himedia-MB083) in PBST and the blots were probed with primary antibodies at 4°C overnight. Following three washes with PBST, the blots were incubated with either anti-rabbit (Invitrogen-65-6120) or anti-mouse (Santa Cruz – SC516102) secondary antibody for an hour. The blots were then washed thrice using PBST, visualized using ECL chemiluminescence detection system (Thermo-34580). Details of the all the primary antibodies used in this study are provided in supplementary information.

### Chromatin immunoprecipitation (ChIP)

ChIP experiments were performed using our laboratory protocol [27, 29] adapted from Fortschegger et al., 2010 with slight modifications [30]. Following the required treatments, cells were washed with PBS, fixed with 5mL of 1% formaldehyde for 10 minutes on a rocker. The reaction was quenched using 1.25M glycine (0.5ml) for 5 minutes at room temperature. Cells were pelleted, washed with PBS and lysed (on ice) for 10 minutes in 300uL of ChIP lysis buffer, followed by sonication (0.3 mm probe sonicator) on ice for 10 minutes on 30s ON-30s OFF cycles at 30% efficiency and diluted using equal amount of ChIP dilution buffer. 100uL of this diluted chromatin was incubated with 1-3ug of primary antibody overnight at 4°C, followed by addition of A/G agarose beads (Santacruz – SC-2003) and agitated for 4hrs at 4°C. The beads with immunoprecipitated complexes were washed 3 times with low salt buffer, once with high salt buffer, once with LiCl buffer and finally with TE buffer. Chromatin elution was performed using fresh elution for 1hr at room temperature, followed by crosslink reversal (using 5M NaCl) at 65°C for 3hrs followed by RNase and proteinase K treatments. The DNA was purified using Qiagen PCR purification kit. Compositions of all the buffers, sequence of the primers used and %INPUT calculations are provided in supplementary information.

### DHE staining

DHE assisted superoxide quantification was performed using confocal microscopy technique. For staining, cells were seeded on a coverslip and treated with KDM5A siRNA and SFB-RBP2 construct for 48hrs, and drug treated (curcumin) for 24hrs. The treated media was decanted, and cells were then washed with PBS, incubated with 5uM DHE (sigma 37291) stain for 30 minutes in the incubator, followed by washing and image acquisition using Leica microscope.

### Evaluation of ROS level using DCFDA by Flow cytometry

Intracellular ROS (peroxide) levels were detected using fluorescent probe DCFDA (sigma D6883) following 24hrs of drug treatment. Cells were treated with curcumin for 24hrs followed by trypsinization, washing with PBS and incubation with 5nM DCFDA for 30 minutes at 37 °C in dark. Cells were then washed and analysed in BD accuriC6 FL1 filter (supported by DST-FIST Level I). Data acquired were analysed using BD software.

### Immunocytochemistry

HeLa and HaCaT cells were seeded on a coverslip in a 6-well plate, transfected with MLL1 siRNA for 48hrs followed by fixation using 100% methanol for 5 minutes, permeabilized with PBST on ice for 10 minutes and blocked using 1% BSA in PBST for 1hr. The cells were incubated with anti-MLL1 antibody overnight at 4°C in a humid chamber. Following 3 washes with PBS, the cells were incubated in Alexa488 (anti-rabbit ab150077) for 1hr at room temperature, washed thrice and images were acquired using Leica microsystems.

### Statistical significance

When 2 groups were tested – Student’s ‘*t*’ test was employed and when multiple classes of data were analysed – either a one-way or two-way ANOVA were employed to test the significance, and the test employed along with the *p*-values and their inference is mentioned in figure legends.

## Results

### KDM5A regulates metabolic landscape in a H3K4me3 demethylation dependent manner

Gene expression profile of KDM5A (demethylase of H3K4me3) and MLL1 (writer methyltransferase of H3K4me3) across multiple tumor samples paired with normal tissues was extracted from UALCAN and gepia2 (**figure 1**), and we noticed that KDM5A expression in cervical tumors (CESC) and metastatic prostate tumors was either marginally higher (**figure 1a-c**) or reduced (as observed in PRAD and gepia2 data) when compared to the normal tissue (**figure 1d**). MLL1 was drastically reduced in both CESC and PRAD patients when compared to their normal counterparts **(supplementary figure S1)**.

**Figure 1.**
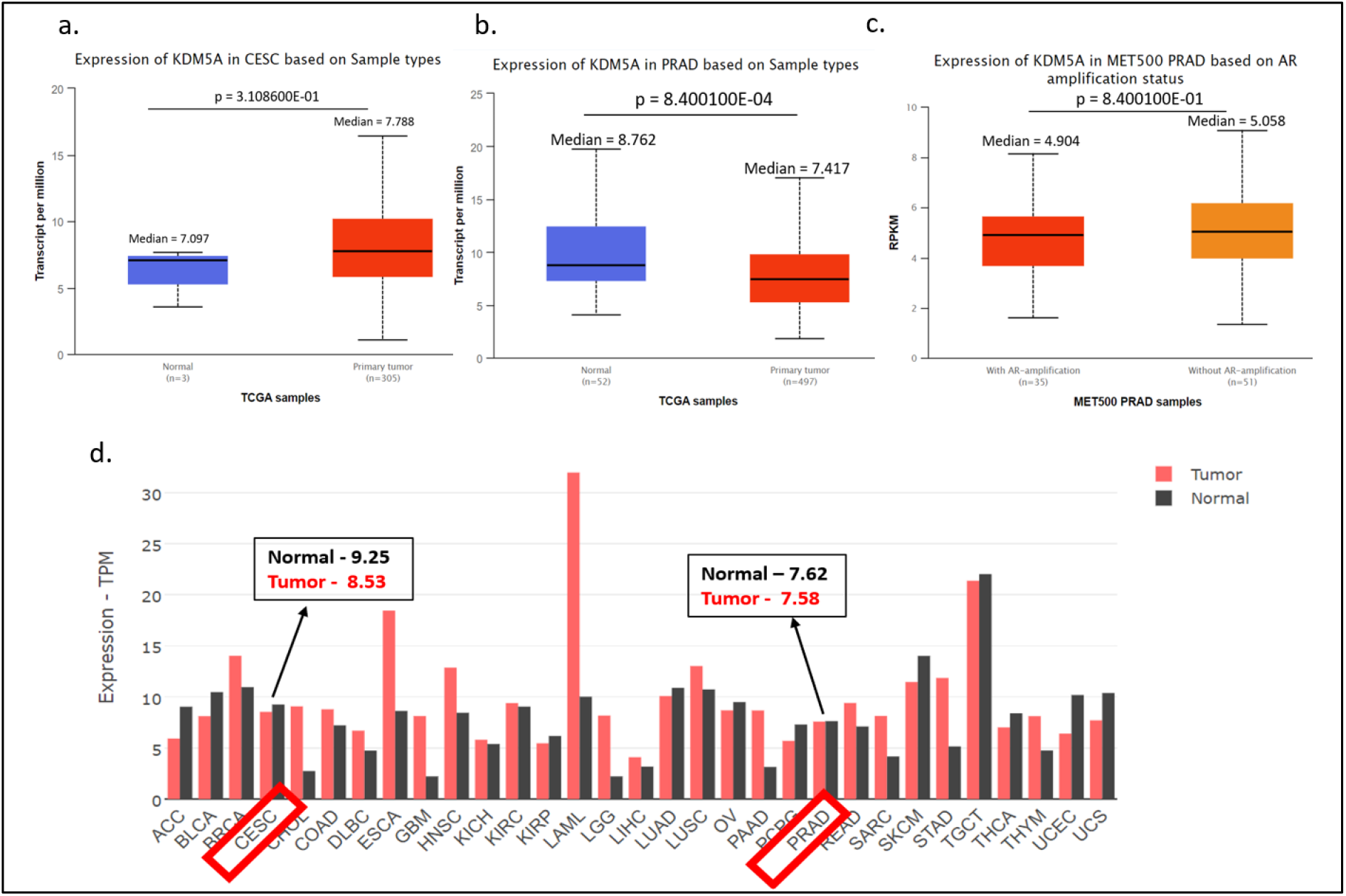
Box plots indicating the expression of KDM5A in cervical squamous cell carcinoma (CESC) and prostate adenocarcinoma (PRAD) tumors compared to their respective normal tissues was obtained using UALCAN database. **(a).** The expression of KDM5A in CESC primary tumors (n=305) was marginally higher than normal tissues (n=3). **(b).** Fold change of KDM5A in prostate cancer primary tumors (n=497) was less compared to normal tissues (n=52) but, **(c).** metastatic prostate cancer tissues (without AR amplification, n= 51) exhibited slightly higher mRNA levels of KDM5A compared to AR-amplification samples (35). **(d).** KDM5A expression profile in CESC and PRAD tumors was less compared to normal tissues as observed from Gepia2 database gene expression profiling data with bars representing median expression of gene in transcripts per million.

Using transcriptional data sets from TCGA, we compared the expression of 80 genes involved in different metabolic processes like glycolysis (19), TCA (19), ATP synthesis (20), PPP (5), ROS scavenging enzymes (13) and genes involved in lactate fermentation (4). From our correlation analysis of comparing expression of these 80 genes from individual patient data sets with KDM5A expression in both CESC **(Supplementary table T1)** and PRAD **(Supplementary table T2)**, we understood that most of the genes (54) exhibited a negative correlation between KDM5A and the positively correlated genes showed marginal scores. We further plotted the comparison of mRNA transcripts of these genes from the two datasets against ranked KDM5A expression and concluded that KDM5A is generally negatively correlated with many metabolically related genes (**figure 2**, **supplementary table T3-T6).** Along with KDM5A, we also extracted MLL1 correlation scores for these genes using the same datasets as above, and besides being a transcriptional activator, MLL1 also showed negative correlation with 55 of the 80 genes (in cervical cancer) and 56 of 80 genes (in prostate cancer) tested **(Supplementary Table T7 and T8)**.

**Figure 2.**
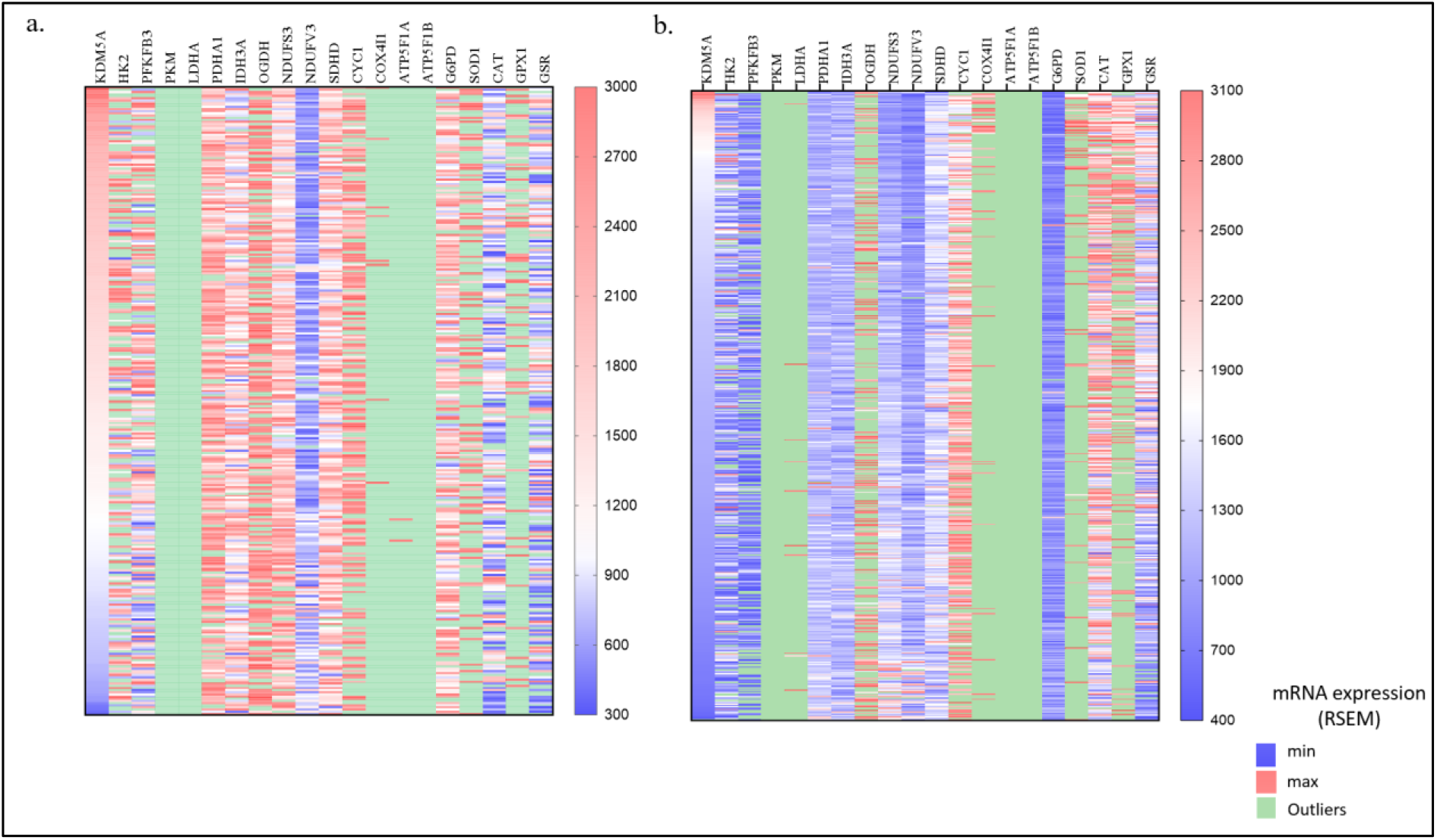
An inverse correlation was observed between genes involved in metabolic processes compared with ranked KDM5A expression in **(a).** cervical squamous cell carcinoma (n=294) and **(b).** prostate adenocarcinoma (n=493). The data represents mRNA expression (RNA-Seq V2 RSEM) from individual patient datasets obtained from TCGA. The individual values and statistical data are submitted as supplementary tables T3-T6.

Analyzing the in-silico studies, we understood that both KDM5A and MLL1 are generally expressed in cervical and prostate cancer patients (slightly enhanced though in metastatic prostate tissue samples). Thus, we sort to elucidate how KDM5A and MLL1 regulate the genes involved in key metabolic processes in these cancers by either depleting (using siKDM5A and siMLL1) or overexpressing (SFB-RBP2) these proteins [29] and **(supplementary figure S2**); we further performed quantitative PCR (23 genes) and immunoblotting (4 proteins) of some of the important enzymes involved in glycolysis, TCA cycle, ATP synthesis, PPP and lactate fermentation using HeLa, HaCaT and PC3 cell lines. From our q-PCR study, we understood that KDM5A regulated most of the genes in a demethylase dependent manner in all the 3 cell lines tested proving that these genes were direct targets, hence displayed increased expression in knockdown and were downregulated when KDM5A was overexpressed. In MLL1 knockdown condition, gene expression was inconsistent among the 3 cell lines, with some genes being upregulated and some others repressed and thus no particular trend could be configured (**figure 3**), which implicates that, there is tissue specificity in MLL1 function.

**Figure 3:**
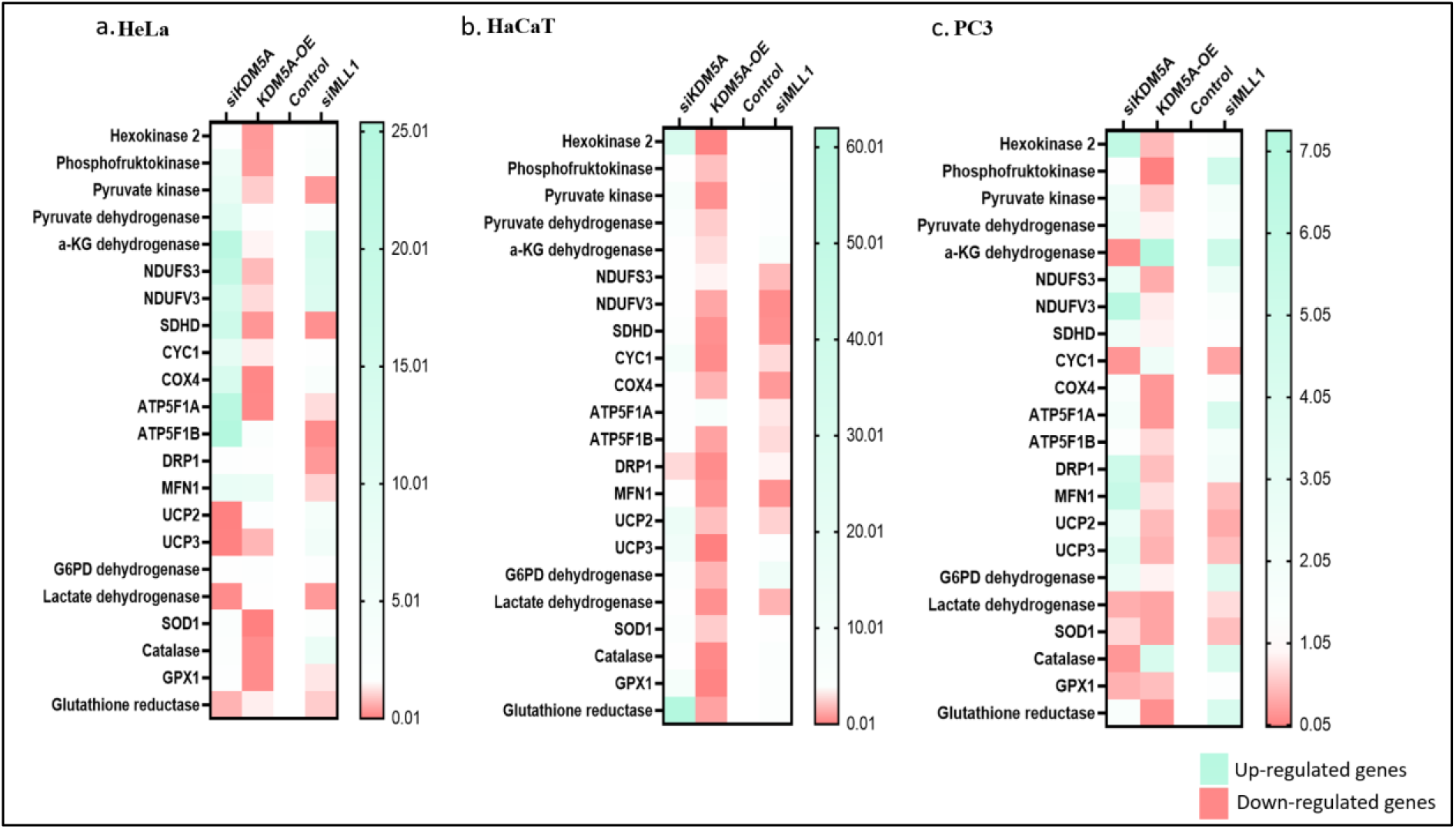
KDM5A and MLL1 regulate the expression of genes related to metabolic processes as analysed by q-RTPCR in HeLa (a), HaCaT (b) and PC3 (c) cell lines treated with siRNA against KDM5A, or MLL1 and transfected with KDM5A overexpression construct, with either negative control siRNA or normal untreated cells used as control, followed by total RNA extraction, cDNA synthesis and quantitative PCR was performed to quantify the indicated gene transcripts (labelled to left of the heatmap). β-actin was used to normalize the mRNA levels, and the fold change is expressed relative to either untreated control or negative control siRNA treated cells (value set to 1). Only one control panel is plotted to simplify the representation. Triplicate sets (n=3) were performed in all 3 cell lines and statistical significance was tested using prism5 software, the data was under p<0.05.

Immunoblotting experiments revealed that the glycolytic enzyme phosphofructokinase (PFK1, the enzyme that catalyses committing reaction of converting F6P to F1,6BP) was regulated in a demethylase dependent manner in HaCaT, but a reverse trend was observed in both the cancerous cell lines (**Figure 4a-b**) but increased in MLL1 knockdown (**figure 4b**) in coherence with the MLL1-PFK gene correlation scores **(supplementary table T7 and T8)**. From correlation analysis (**figure 4a**) it is clear that IDH3A exhibited negative correlation with both KDM5A and MLL1 **(supplementary table T7 and T8)**, and the western blots replicated the same in all the 3 cell lines tested, i.e., IDH3A enhanced in siKDM5A treatment (knockdown of KDM5A), and reduced in KDM5A overexpression; however, in modulation of MLL1, IDH3A either enhanced slightly (HaCaT and PC3) or remained same (HeLa). Of the 20 ATP synthesis genes studied in our correlation analysis **(Supplementary table T1 and T2)**, 18 genes exhibited negative correlation with KDM5A, of which ATP synthase subunit (ATP1F1) was immunoblotted and we configured that it was a direct target of KDM5A but remained largely unregulated in MLL1 knockdown (**figure 4b-c**). Taken together, among all the genes tested here, TCA cycle and ATP synthesis related genes are direct targets of KDM5A enzymatic function in both cervical and prostate cancer cell lines along with the normal keratinocyte (HaCaT), excluding the glycolytic enzyme encoding gene.

**Figure 4:**
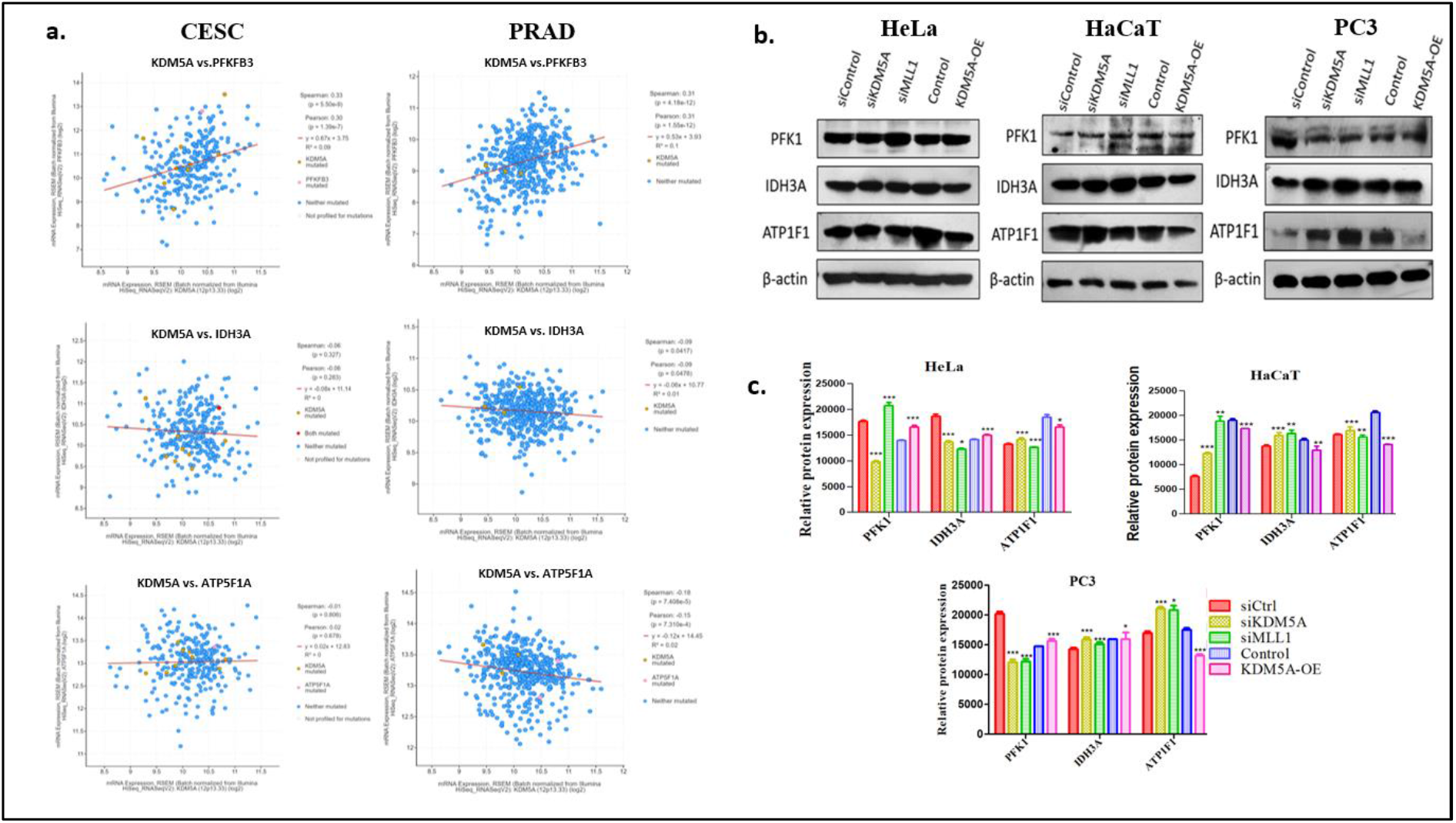
In cervical and prostate cancers: KDM5A regulates TCA cycle and ATP synthesis directly in a demethylase dependent manner. **(a)**. Scatter plots showing correlation of KDM5A with PFKFB3, IDH3A and ATP1F1A using RNA-seq datasets of cervical squamous cell carcinoma (CESC) and prostate adenocarcinoma (PRAD) patients from TCGA. PFKFB3 showed a positive correlation score, and all the other genes are negatively correlated with KDM5A in both cancer patient samples. **(b)**. Western blotting showing expression of PFK1, IDH3a and ATP1F1 in HeLa, HaCaT and PC3 cell lines following treatment of cells with siRNA against KDM5A and MLL1 and KDM5A overexpression conditions. Specific antibodies used to probe the blots are indicated to the left of each panel. Immunoblots were performed in triplicate (n=3), and mean fluorescence measured through ImageJ tool was plotted into bar graphs with SE are presented along the blots **(c)**.

As most of the cancerous cells also use pentose phosphate pathway (PPP) as an alternative source of generating precursors of biomolecules, we analysed how KDM5A regulates this metabolic derailing in both the cancers (HeLa and PC3). In cervical patient samples, 4 of the 5 genes showed negative correlation (with correlation score ranging from -0.015 to -0.3) **(Supplementary table 1** and **figure 5a**) and in prostate adenocarcinoma, all 5 genes were negatively correlated (higher scores compared to CESC i.e., range of -0.1 to -0.6) **(Supplementary table 2** and **figure 5a**). We further immunoblotted G6PD (the rate limiting enzyme of PPP) and understood that in HaCaT and PC3, KDM5A acts as a demethylase, but in HeLa, the expression of G6PD was largely unregulated in all treatments (**figure 5b, c**). This is consistent with the low correlation coefficient between KDM5A vs G6PD in CESC (Spearman coefficient = -0.015). G6PD was known to be the key to malignancy in cervical cancer as a very strong signal of G6PD was observed in all cervical cancer patients screened compared to prostate cancer **(supplementary figure S3)**, and its suppression inhibited migration capabilities of HeLa cells [31]. In MLL1 knockdown, we observed an increase in G6PD expression in all three cell lines, which was consistent with the correlation analysis **(supplementary table T7 and T8)**. Further, to confirm the demethylase activity on G6PD promoter, we used HaCaT chromatin and immunoprecipitated with H3K4me3, and observed that in siKDM5A – an increase in H3k4me3 occupancy was observed, and KDM5A overexpression, a decrease was noted (**figure 5d**).

**Figure 5:**
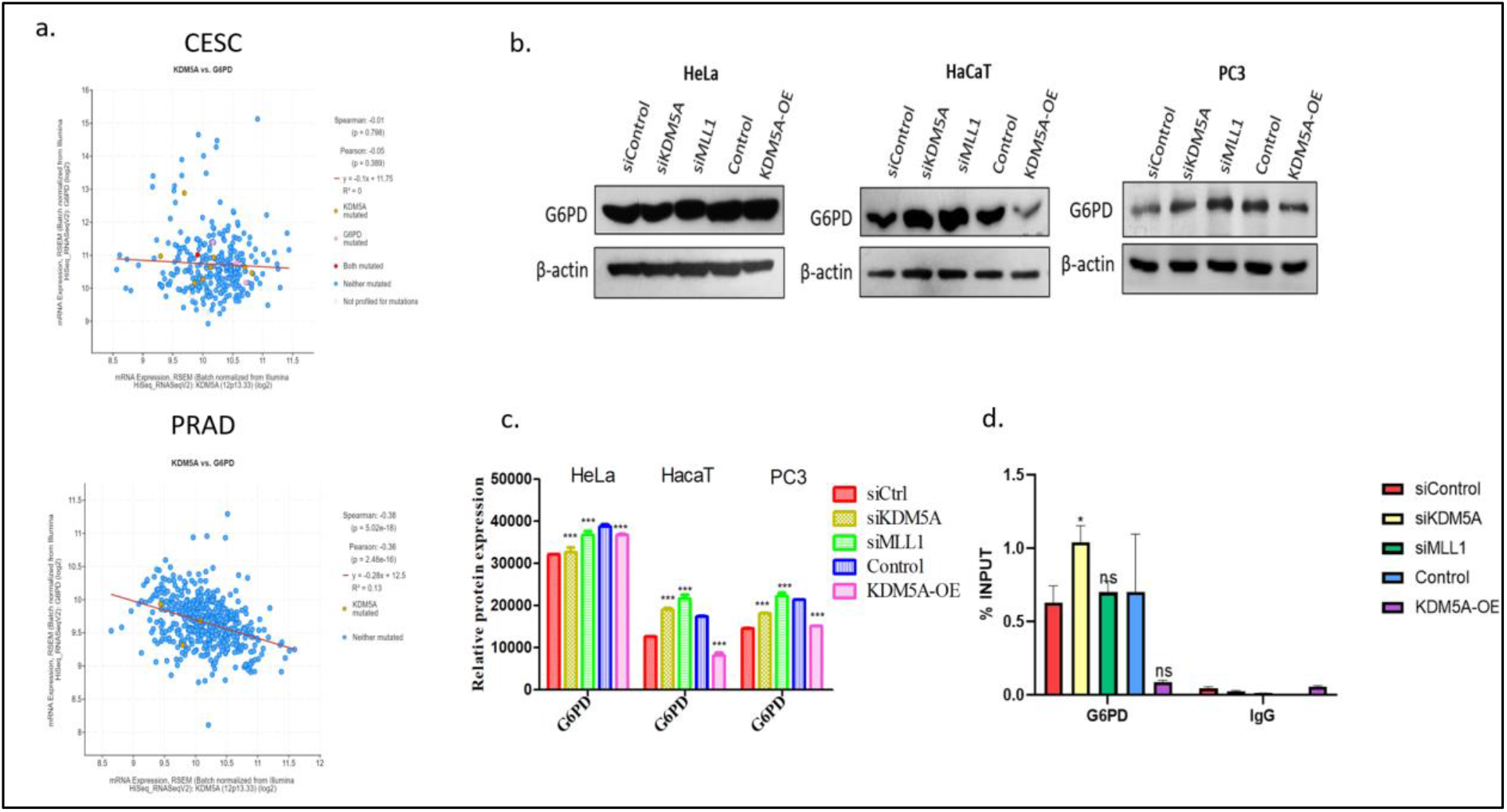
KDM5A shows tissue specificity when monitoring pentose phosphate pathway. **(a).** Correlation between KDM5A and G6PD in CESC and PRAD is depicted as a scatter plot where marginal correlation observed in CESC and a negative correlation with spearman coefficient of -0.38 in PRAD was observed. **(b).** Immunoblots with G6PD expression: No change was observed in HeLa, but in HaCaT and PC3, KDM5A acts as a demethylase on this promoter. Immunoblots were performed in triplicate (n=3), and mean fluorescence measured through ImageJ tool was plotted into bar graphs with SE are presented along the blots **(c). (d).** ChIP data following siRNA against KDM5A and MLL1 and overexpressing KDM5A in HaCaT cells shows that KDM5A regulates G6PD promoter in a demethylase dependent manner; this experiment was performed in duplicates (n=2), and the error bars represent SD, Bonferroni test was used to calculate the significance. *P≤0.05, **P≤0.01, ***P≤0.001 and ns is non-significant i.e., P>0.05.

To understand the impact of KDM5A on the overall structure of mitochondrial networking, we estimated the mitochondrial fission (DRP1) and fusion (MFN2) proteins. Although in PC3, KDM5A act as a classical repressor on MFN2, but in HeLa and HaCaT cell lines, we observed high levels of MFN2 in all the treatments. With respect to KDM5A, the fission protein DRP1 largely remained unaffected in the cancer cell lines, but in HaCaT, a slight regulation was observed which was consistent with KDM5A being a demethylase. Thus, it is difficult to draw a general trend in the regulation of mitochondrial structure by KDM5A, but we understand that rate of proliferation (in case of MFN2) and tissue specificity (in case of DRP1) affected the regulation pattern. Regarding MLL1 control over these genes, it displayed a negative correlation to MFN2, but no particular regulation tendency was observed on DRP1 (although a slight increase in cancer cells and decrease in normal cell line was observed, the regulation was inconsistent) (**figure 6a, b**).

**Figure 6:**
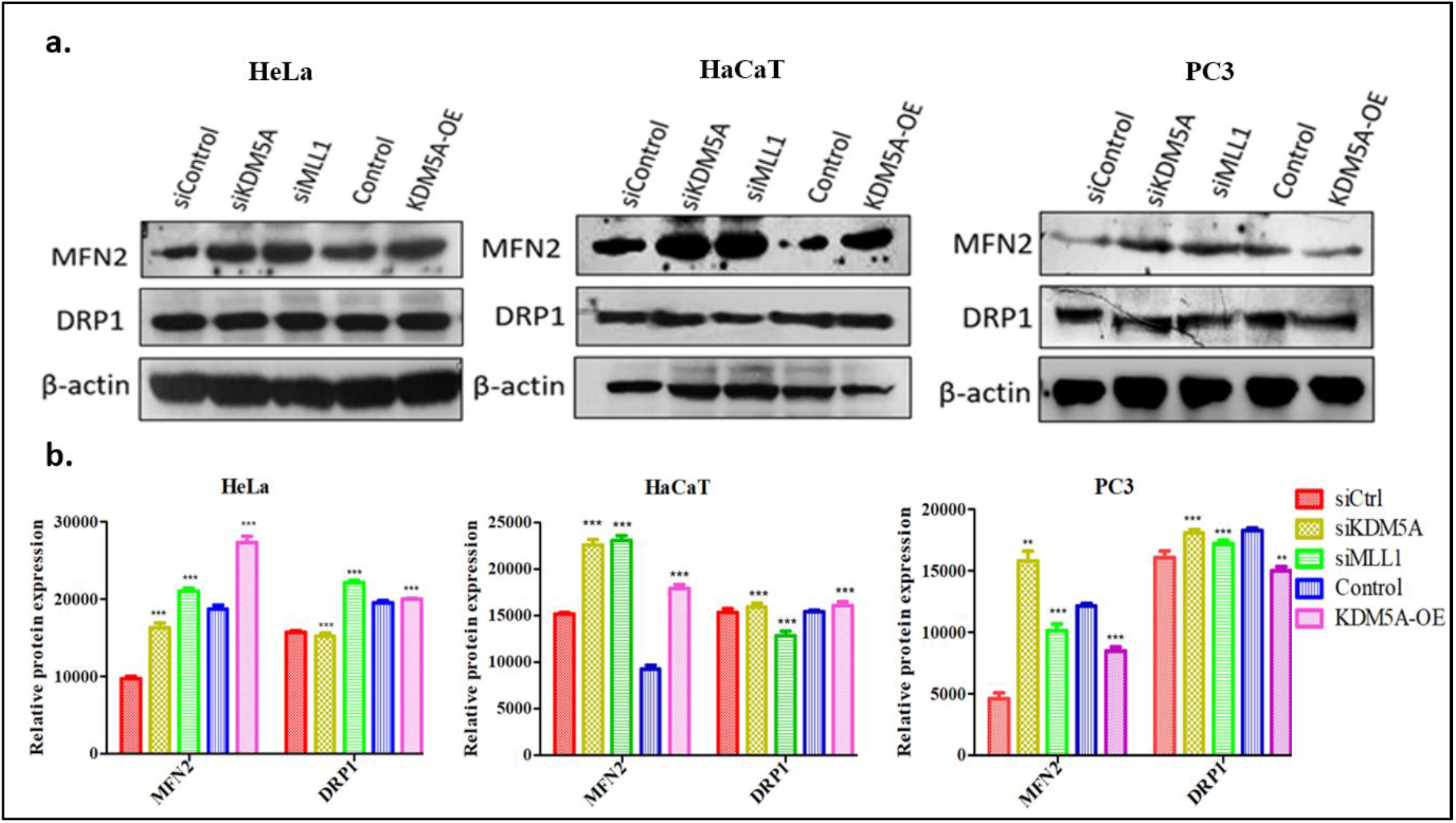
**(a)**. Regulation of mitochondrial mass by KDM5A and MLL1. Although KDM5A acts as a demethylase tissue specifically (in PC3) on MFN2 promoter, no particular correlation could be drawn from the immunoblots in case of DRP1. The western blots were performed in triplicates and bar graphs representing mean with SE are represented along with the data **(b)**.

### KDM5A regulates cellular ROS level

Redox status defines the proliferative and viability index of cells by controlling different signalling pathways involved in cell survival, apoptosis and division and ROS scavenging enzymes play a crucial role in the maintenance of a homeostasis, but many cancers are said to have altered levels of ROS leading to oxidative stress. Our correlation analysis showed that out of the 13 genes tested against KDM5A, 8 genes in CESC and 5 genes in PRAD had negative scores **(Supplementary table 1 and 2)**, and it was surprising to note that while SOD group enzymes were negatively correlated with KDM5A, but Catalase has a positive relation (**Figure 7a**). Thus, we analysed the regulation of SOD1 and catalase by KDM5A and MLL1. Although both enzymes were regulated in demethylase dependent manner, but catalase remained unregulated in both cancer cells. Unlike the other genes of glycolysis and TCA cycle which were upregulated in siMLL1 condition, MLL1 suppression did not enhance the expression of this class of genes (except catalase in HaCaT) (**Figure 7b-c**). We further quantified the superoxide levels in HeLa using dihydroethidium (DHE) staining, which oxidizes in the presence of superoxide to form ethidium that intercalates with DNA. In agreement with our western data, ethidium staining in siKDM5A treated cells decreased (**figure 8a, c**) as SOD1 increased and intense staining was detected in KDM5A overexpression as SOD1 levels fall (**figure 8b, c**).

**Figure 7:**
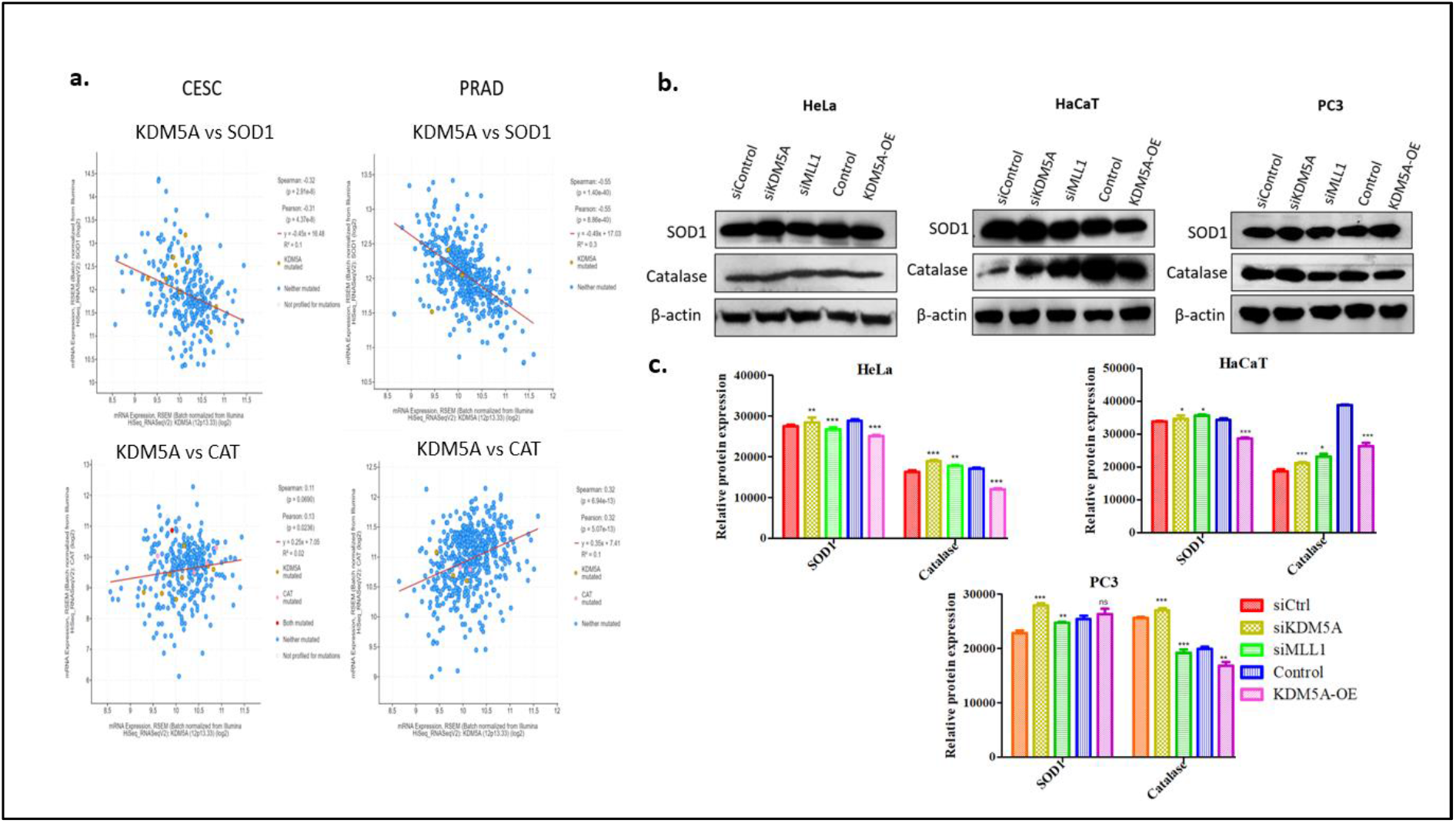
Regulation of ROS scavengers by KDM5A and MLL1**. (a).** KDM5A has a negative correlation with SOD1 but a positive correlation with catalase in both CESC and PRAD **(b).** KDM5A regulates SOD1 by demethylating H3K4me3 but doesn’t act similarly on catalase promoter in cancer cells. Immunoblots were performed in triplicate (n=3), and mean fluorescence measured through ImageJ tool was plotted into bar graphs with SE are presented along the blots **(c)**.

**Figure 8:**
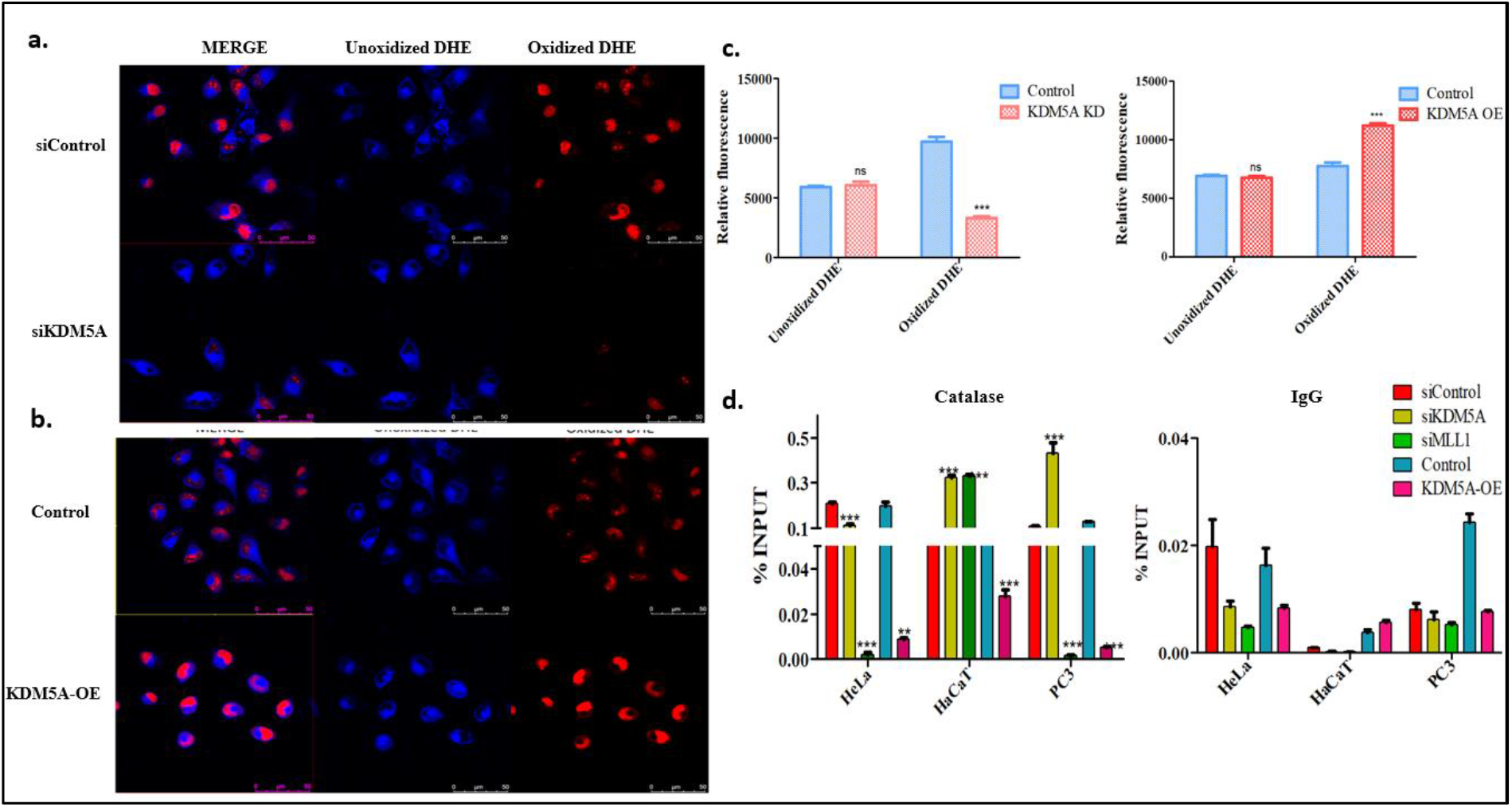
Quantifying ROS levels and mechanism of gene regulation **(a-b).** Superoxide quantification (in HeLa) using DHE staining reveals reduced ethidium intercalating DNA in siKDM5A as SOD1 levels increased and vice versa in KDM5A overexpression, the florescence intensity from triplicates was quantified by ImageJ and presented as bar graphs with mean+ SEM. Statistical significance was tested using prism5 **(c). (d)** ChIP data following siRNA against KDM5A and MLL1 and overexpressing KDM5A, to study the occupancy of H3K4me3 revealed that in HaCaT, promoter H3K4me3 status dictates catalase expression but in cancerous cells (HeLa and PC3) other factors also contribute to the overall protein levels. This experiment was performed in duplicates (n=2) **(d).** IgG ChIP data, the error bars represent SD, Bonferroni test was used to calculate the significance. *P≤0.05, **P≤0.01, ***P≤0.001 and ns is non-significant i.e., P>0.05.

Further, we screened ChIP-seq data generated using KDM5A (SRX1721564) and H3K4me3 (SRX1721545) as antigens in T-47D and MLL1(SRX4931256) and MLL2 (SRX2007818) in MCF7 to understand their occupancy and regulation of metabolic genes. We noted that MLL2 overlapped with both H3K4me3 and KDM5A binding regions **(supplementary figure 4)**. We then performed chromatin immunoprecipitation using H3K4me3 on catalase promoter (**figure 8d**) to understand its differential regulation in cancer versus normal cells. The active mark increased in siKDM5A but reduced in both siMLL1 and KDM5A overexpression in HaCaT and PC3 cells but was reduced in all 3 conditions in HeLa. Despite the similarity in H3K4me3 promoter status in PC3 and HaCaT catalase gene, the protein expression in PC3 was inconsistent, thus raising questions regarding the translational regulation.

Concluding so far, we traced that both MLL1 and KDM5A are downregulated in cervical and prostate cancers, and that KDM5A, in general, acts in a demethylase dependent manner on most of the metabolic genes, with tissue specific regulation observed in case of G6PD and catalase genes. MLL1 on the other hand does not activate these processes as knockdown of MLL1 did not affect their expression and instead an enhanced expression with increased H3K4me3 was noted, which could be a function of compensation from other methyltransferases (like MLL2 and SET proteins).

### Curcumin mediated suppression of metabolism involves KDM5A

We have used phytochemicals of different classes to test their effect on regulating KDM5A and MLL2, and their ineffectiveness towards normal cells. We have utilized a monoterpene (geraniol), a benzoquinone alkaloid (embelin), a phenolic acid (gallic acid), a triterpene (stigmasterol), and a polyphenol (curcumin) compound. All these compounds are naturally available from edible sources (**Table 1**).

**Table 1:** Classification of phytochemicals used in this study along with their naturally available sources.

| Phytochemical compound | Class | Natural sources |
| --- | --- | --- |
| Geraniol | Terpene | Many essential oils of fruits, vegetables, and herbs including rose oil, citronella, lemongrass, lavender, and other aromatic plants |
| Embelin | Benzoquinone alkaloid | Embelia ribes, Aegiceras corniculatum, Embelia schimperi, Erythrorhiza, Lysimachia punctata |
| Gallic acid | Polyphenol | Blueberry, blackberry, strawberry, plums, grapes, mango, cashew nut, hazelnut, walnut, tea, wine |
| Stigmasterol | Phytosterol | Fats and oils of soybean, calabar bean and rape seed, vegetables, legumes, nuts, seeds, and unpasteurized milk |
| Curcumin | Polyphenol | Curcuma longa |

Following cell viability assay **(supplementary figure S5)**, we treated cells with the phytochemical compounds for 24hrs followed by immunoblotting for KDM5A and MLL2, and detected that in normal cells (HaCaT), KDM5A remained unchanged but in HeLa and PC3 cells, GA, stigmasterol and curcumin treatments enhanced KDM5A and MLL2 levels either reduced slightly or remained unaltered (**figure 9a-b**). As curcumin showed consistent regulation in both HeLa and PC3 i.e., enhanced KDM5A and reduced MLL2 and a counter regulation in HaCaT, we tested the global changes in H3K4me3 following curcumin treatment and observed that in cancerous cells (HeLa), the active mark reduced (due to increased KDM5A and reduced MLL2) (**figure 9c**) but in normal cells the H3K4me3 status slightly increased (as KDM5A is unchanged and MLL2 is enhanced) (**figure 9c**).

**Figure 9:**
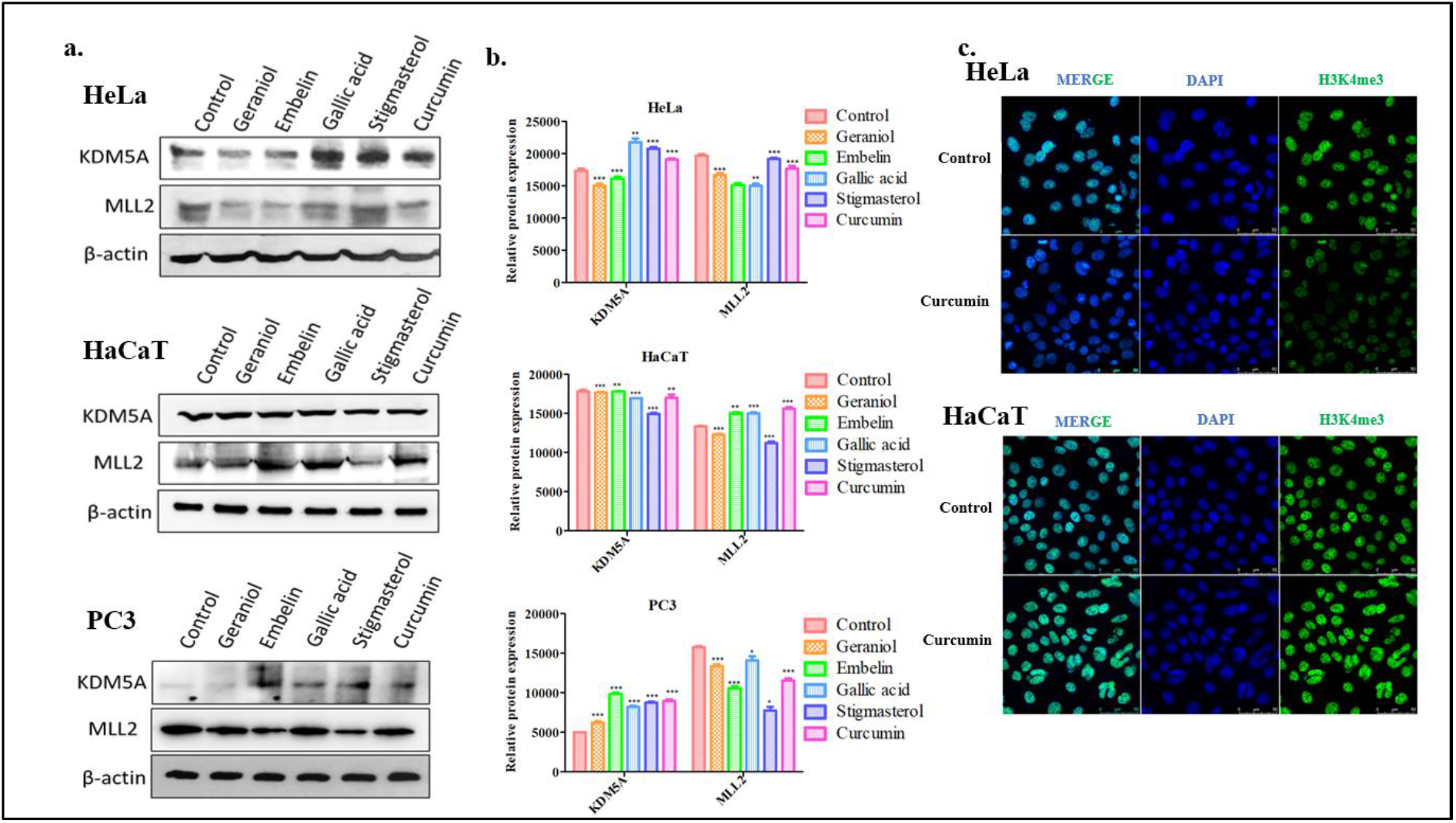
Effect of different phytochemical compounds on the expression of KDM5A, MLL2 and global changes in H3K4me3. **(a).** Western blots showing the expression KDM5A and MLL2 following treatment with Geraniol, embelin, GA, stigmasterol and curcumin. Immunoblots were performed in triplicate (n=3), and mean fluorescence measured through ImageJ tool was plotted into bar graphs with SE are presented along the blots **(b)**. **(c).** Immunocytochemistry against H3K4me3 following treatment with curcumin in HeLa and HaCaT cells respectively.

We then tested if the compounds had an inhibitory effect on the enzymes related to glycolysis, mitochondrial respiration, PPP, and mitochondrial fission. From our immunoblotting data (**Figure 10a-b**), we noted that except in curcumin treatment, the expression of metabolic signature genes (IDH3A, ATP1F1, G6PD) and mitochondrial fission gene (DRP1) remained unaltered or slightly enhanced in other drugs. In curcumin treatment, HeLa exhibited a decline in all the 4 genes tested and in PC3 – IDH3A and ATP1F1 were reduced.

**Figure 10:**
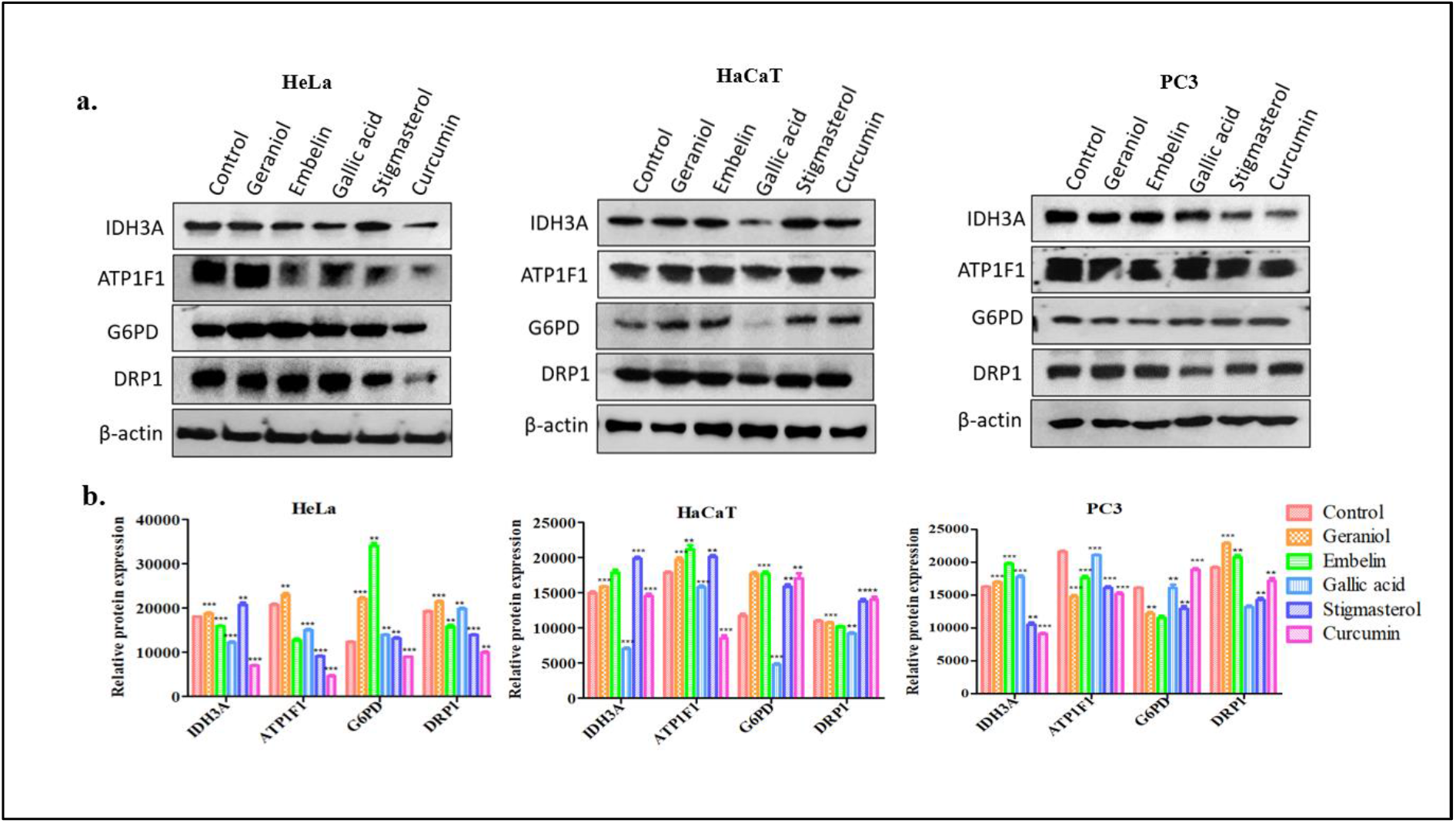
Profiling metabolic signature genes by immunoblotting **(a)** in HeLa, HaCaT and PC3 following treatment with phytochemical compounds like Geraniol, embelin, GA, stigmasterol and curcumin. Curcumin treatment inhibited mitochondrial function in cancerous cell lines but was ineffective towards normal cells. Immunoblots were performed in triplicate (n=3), and mean fluorescence measured through ImageJ tool was plotted into bar graphs with SE are presented along the blots **(b)**.

As observed earlier, curcumin treatment in HeLa increased KDM5A expression, so we further wanted to test whether the reduction in metabolic genes (IDH3A, ATP1A1 and G6PD) and the fission protein DRP1 was due to enhanced KDM5A expression following curcumin treatment, so we performed transient knockdown of KDM5A in curcumin treated HeLa cells. From our immunoblots (**Figure 11**), we observed that together curcumin and siKDM5A treatment led to restoration of all the proteins to normal levels when compared to curcumin treatment alone (where they were reduced), validating that curcumin treatment mediated decrease in mitochondrial activity in HeLa cell line was KDM5A dependent and reducing KDM5A could restore the expression of metabolic genes to normality.

**Figure 11:**
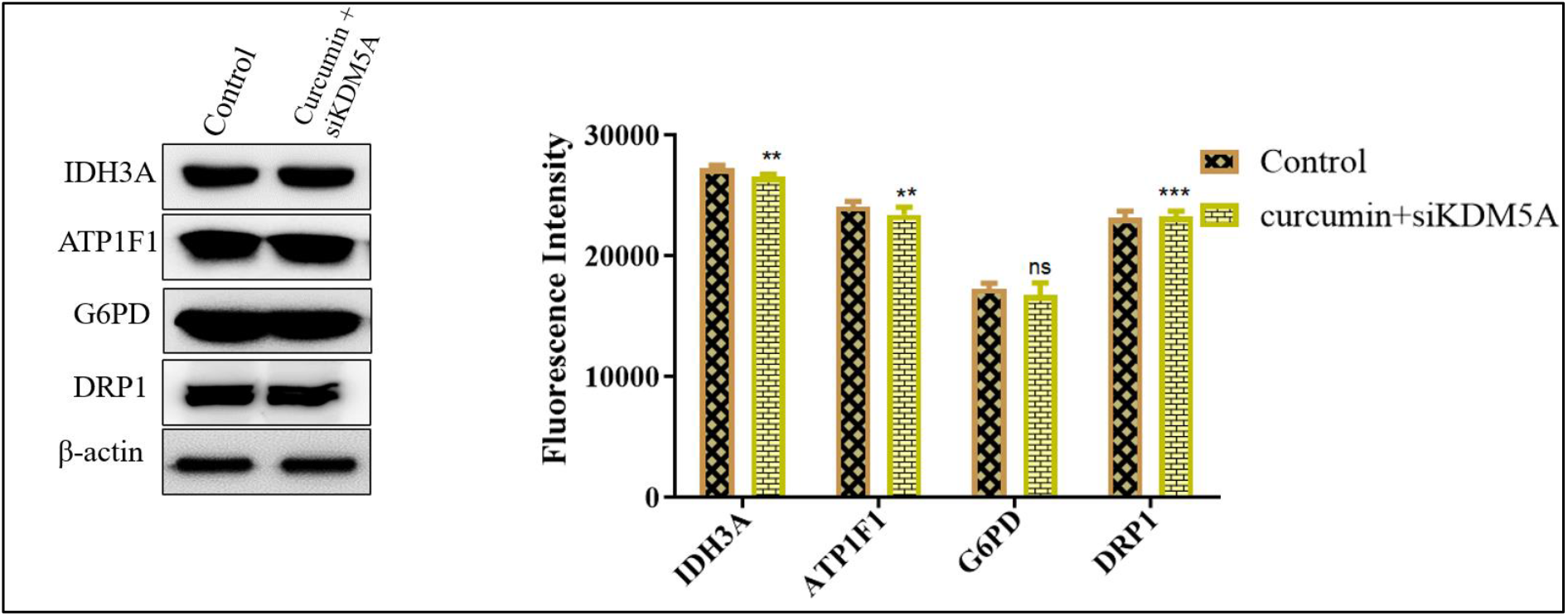
Knockdown of KDM5A in curcumin treated HeLa cells led to restoration of metabolic gene expression, which was reduced in curcumin treatment alone (due to enhanced KDM5A expression.

Further, ROS scavengers SOD1 and Catalase were quantified, and curcumin treatment showed differential expression of SOD1 in Hela (slightly enhanced) and PC3 (reduced), and catalase expression reduced in both the cancerous cell lines when compared to HaCaT (unregulated) (**figure 12a-b**). DHE staining in curcumin treated HeLa cells revealed reduced ethidium-stained DNA compared to control (**figure 12c-d**) and DCFDA quantification for peroxide levels showed enhanced H2O2 in curcumin treated HeLa (where catalase levels were reduced) when compared to HaCaT (unchanged) **(figure 12e)**. As MLL1 showed a negative correlation with most of the metabolic genes, we tested MLL2 occupancy as it is the principal methyltransferase at genes involved in housekeeping functions. Our ChIP data revealed reduced H3K4me3 as a function of enhanced KDM5A and reduced MLL2 on Catalase promoter in HeLa cells but an enhanced H3K4me3 with high KDM5A and MLL2 occupancy in HaCaT cells **(figure 12f)**. From this, we understood that the regulation of metabolic genes is by H3K4me3 is monitored by the KDM5A/MLL2 ratio.

**Figure 12:**
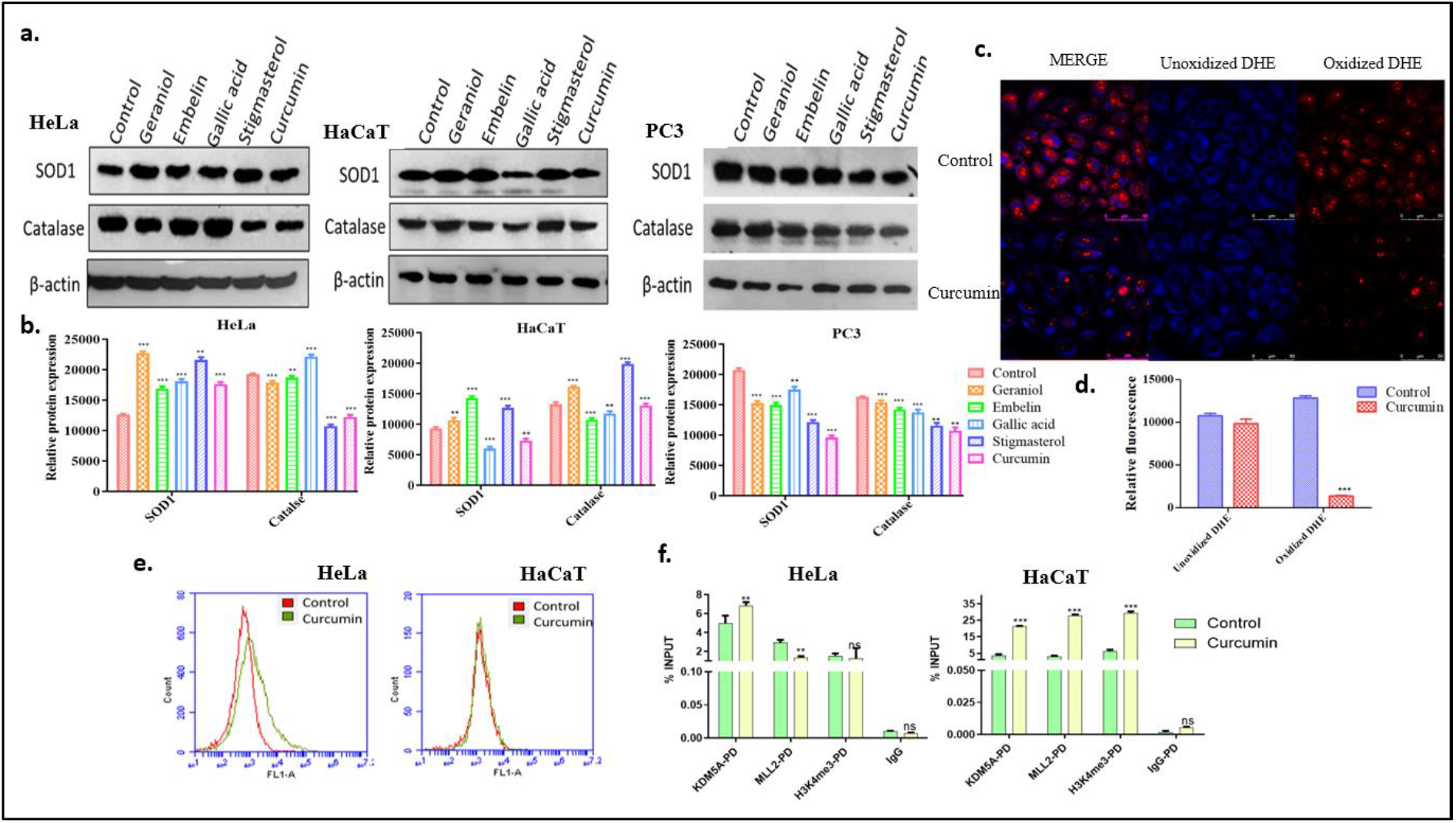
Curcumin regulates ROS levels in HeLa and PC3 cells. **(a).** Drug treated HeLa, HaCaT and PC3 whole cell lysates were immunoblotted with SOD1 and catalase antibodies. Immunoblots were performed in triplicate (n=3), and mean fluorescence measured through ImageJ tool was plotted into bar graphs with SE are presented along the blots **(b). (c)**Superoxide levels in curcumin treated were quantified using DHE stain, untreated cells were used as control, the florescence intensity from triplicates was quantified by ImageJ and presented as bar graphs with mean+ SEM. Statistical significance was tested using prism5 **(d)**. **(e).** DCFDA stained HeLa and HaCaT cells were subjected to flow cytometry, to study changes in H2O2 levels following curcumin treatment. **(f).** KDM5A, MLL2 and H3K4me3 occupancy on catalase promoter following curcumin treatment inn HeLa and HaCaT cells. ChIP was performed in duplicates (n=2). The error bars represent SD, Bonferroni test was used to calculate the significance. *P≤0.05, **P≤0.01, ***P≤0.001 and ns is non-significant i.e., P>0.05.

To further assess whether enhanced ROS in HeLa cells following curcumin treatment led to apoptotic cell death, we performed annexin-V staining. From **figure 13**, we could confirm that curcumin treatment could increase apoptotic population compared to control in HeLa cells. To test if curcumin induced ROS was responsible for the apoptosis, we quenched the ROS using N-acetylcysteine (NAC) to see if any reduction in apoptotic population could be observed. As expected, curcumin with NAC treatment led to a reduction in apoptosis, making it clear that curcumin mediated increase in KDM5A reduced ROS scavenging enzymes thereby accumulating free radicles, which increased apoptotic cell death in HeLa cells, and quenching the ROS species using NAC decreased the extent of apoptotic cell death. In HaCaT cells, as anticipated, no changes in apoptosis were detected between control and curcumin treated cells, which is reasonable as curcumin treatment does not alter either KDM5A or ROS levels in HaCaT cells.

**Figure 13:**
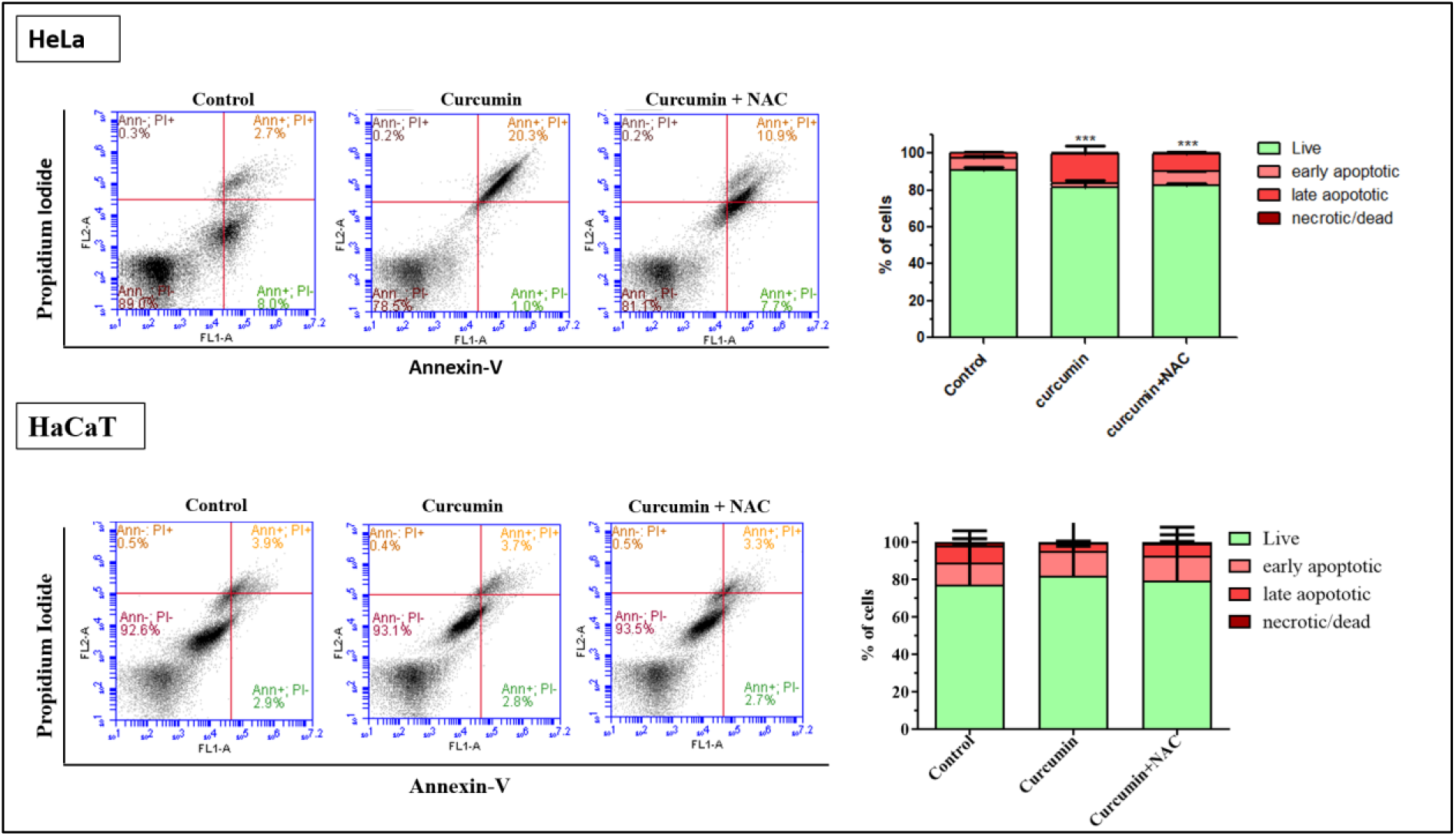
HeLa cells treated with curcumin or curcumin+ NAC and stained with annexin-V to study apoptosis using flow cytometry. Curcumin treatment enhanced apoptotic population in HeLa, which was reduced following quenching of ROS by NAC.

## Discussion

Metabolic alterations are a major hallmark of cancer and adaptation to a certain metabolic phenotype is determined by many factors i.e., mutations in metabolism related genes, proliferative potential of cells, tumor microenvironment, the redox status of the tumor and availability of nutrients. Switching between different phenotypes (metabolic plasticity) involves shutting down a class of genes and activating genes that can help cells better adapt to the microenvironment. This shift is attained in part by recruiting epigenetic modifiers to the gene promoters to either activate or repress downstream targets. One such modification mark is H3K4me3, which is catalyzed by MLLs/SET domain containing proteins and erased by KDM5 family of demethylases.

As cancer cells depend on aerobic glycolysis, enzymes of this pathway are upregulated in many cancers. The key glycolysis rate-limiting enzymes hexokinase-II (HK-II), phosphofructokinase (PFKFB3), and pyruvate kinase M2 (PKM2) along with lactate dehydrogenase (LDHA) were considered for study as these enzymes catalyse irreversible committing reactions and are generally associated with proliferative phenotype [32]. TCA cycle provides energy and macromolecules to the cells and many enzymes like IDH and SDHD are dysregulated in cancers [33], so we analysed pyruvate dehydrogenase (PDH), Isocitrate dehydrogenase (IDH3A) and α-KG dehydrogenase (α-KGDH-E1). Complex-I enzymes NDUFS3 (which initiates the assembly of complex and is essential for catalytic activity) and NDUFV3 (minimal assembly unit required for catalytic activity); Complex-II SDHD; Complex-III CYC1; Complex-IV COX4 and Complex-V ATP5F1A and ATP5F1B were also studied. As altered KDM5A led to morphological changes in mitochondria (either fragmented or elongated – in SAOS-2 cells [22]), we choose to study the fission (DRP1) and fusion (MFN1&2) proteins. To understand if the total ATP output is attained or the proton gradient is dissipated by uncouplers, we also estimated UCP2 and UCP3 as they decrease membrane potential of mitochondria. The gatekeeper or the enzyme that catalyzes the rate limiting step of PPP – Glucose 6-phosphate dehydrogenase (G6PD) was chosen. We have extracted TCGA mRNA expression status of the above-mentioned genes and compared their relation to KDM5A’s levels in cervical and prostate cancer patients.

Here we show how KDM5A generally acts as a demethylase on most of the metabolic gene promoters, as both siKMD5A and SFB-KDM5A (overexpression) treatments exhibited a negative correlation with metabolic gene activity. But in case of MLL1 (a known activator of transcription), contrasting results were observed in siMLL1 treatment, as theoretically, its suppression should ideally downregulate the metabolic genes, instead we observed an increase in the expression of all the genes tested, which could be due to compensation by other activators (MLL2, SET proteins or HATs), as suppression of MLL1 did not lead to reduction of promoter H3K4me3 on catalase gene in HaCaT cell line. Though KDM5A was a transcriptional repressor on most of these genes tested here, tissue specificity was observed in KDM5A vs G6PD in HeLa.

As KDM5 mutant flies show enhanced levels of superoxide [23–24] and KDM5A/B overexpressing tumors also exhibit increased H2O2 levels, we estimated the expression of the cellular antioxidant system, that scavenge ROS, i.e., superoxide dismutase (SOD1)-which converts superoxide anions to H2O2 [5], catalase – reduces H2O2 to water and oxygen [34], glutathione peroxidase 1 (Gpx1) – eliminate H2O2 [35] and glutathione reductase – maintain reduced glutathione levels. Unlike Drosophila KDM5, human KDM5A regulated ROS related genes (SOD1 in all 3 cell lines and catalase in HaCaT) in a demethylase dependent manner, but catalase gene remained unaffected to KDM5A alteration in both the cancerous cells i.e., HeLa and PC3.

As KDM5A levels remained low in cervical and prostate cancer, which could lead to enhanced metabolic activation, allowing an increased glycolytic flux through TCA and PPP, and enhancing the survival and progression of tumors as cervical cancers thrive on PPP [31]. Glycolysis is the key source of energy in prostate cancers due to defective mitochondrial OXPHOS complex activity [36], and thus many glycolytic enzymes were observed to be upregulated and are prognostic markers for prostate cancer detection [2, 37]. Also, an enhanced H3K4me3 status was reported in metastatic prostate cancer patients [38]. These metabolic changes provide energy and biomass to the tumor directing it to a more aggressive phenotype, we hypothesized that an increase in KDM5A would reduce the metabolic rate and increase the oxidative stress (as KDM5A also downregulates SOD1) which could lead to deprivation of biomass and induction of ROS mediated apoptosis in cancers.

Also, as reduced KDM5A enhance SOD1 levels which help cancer cells to better adapt to the increasing oxidative stress by inhibiting ROS mediated activation of apoptotic pathways, this could be an alternative survival route adapted by cervical and prostate cancers. Taken together, the reduced KDM5A phenotype observed in cervical and prostate cancers might supports the cells tumorigenic capabilities by limiting ROS accumulation produced by glycolytic and TCA cycle by increasing scavenging enzymes. Thus, if any compound that can enhance the expression of KDM5A would be a double hit by not only reducing the metabolic rate but also increase the oxidative load in these cancers leading them to energy deprivation and ROS mediated apoptotic cell death. So, we tested some phytochemical compounds [39–40] to see if an increase in KDM5A levels could be obtained thereby hindering metabolic activities in cancer cells. We have chosen a compound from each class of phytochemicals like terpene - geraniol [41–42], alkaloid – embelin [43], phenolic acid - gallic acid [44–46], a phytosterol i.e., stigmasterol [47–49] and polyphenol – curcumin [50–55]. These phytochemicals, when taken as dietary supplements, can modulate gene expression through regulating epigenetic mechanisms involving DNMT, HMT or HDAC activity [56–58]. Treatment of bone marrow derived macrophage (BMDM) cells and LNCap (prostate cancer cell line) with curcumin led to reduction of H3K4me3 methylation [59, 60]. Herein, we also report that, although GA, stigmasterol and curcumin could enhance KDM5A, GA affected the metabolic activity in normal cells (HaCaT) as expression of genes like IDH3A, G6PD, DRP1, SOD1 and catalase was reduced (figure 10 and 12a) and stigmasterol did not show very prominent effect on metabolic genes in cancers as compared to curcumin, which exhibited the best outcome in both cancers by reducing metabolism and enhancing ROS levels.

In our early studies, in quest of epigenetic basis of prostate cancer, we found that DNA methyltransferases and histone deacetylases are highly expressed and controls a group of genes involved in cell cycle regulation and tumor suppression [61–64]. Recently, we have analyzed the roles of epigenetic signaling [65], deciphered the impact of Hh signaling [66], effect of phytochemicals [28], contributions of pluripotency inducing factors [67–69] and the roles of miRNAs in the regulation of prostate cancers [70, 71]. Epigenetic modulations of genes in cancer cell metabolisms are reviewed recently [72–74] All these data from ours and others implies that epigenetic modulations due to varied/aberrant signaling, varied metabolism and cellular microenvironment, and tissue specificity vis-à-vis regulation of genes (involved in those processes) by epigenetic mechanisms play crucial roles in the control of prostate and other cancers.

## Supporting information

Yes

Yes

## Acknowledgements

R. Kirtana received fellowship from CSIR, Govt. of India (CSIR-09/983(0018)/2017-EMR-I). S. Manna is thankful to NIT-Rourkela for fellowships under the Institute Research Scheme, NIT-Rourkela. We thank Dr. Shweta Tyagi, CDFD Hyderabad for providing SFB-RBP2.

## Author Contributions

SKP conceptualized the project. KR and SM performed all the experiments. SKP, KR and SM analysed the data. KR and SM prepared the figures. KR wrote the draft manuscript and SKP edited the manuscript to its final version.

## Funding

This work is supported in part by the Department of Biotechnology (Government of India) project No.: BT/PR21318/MED/12/742/2016 to SKP and by a special research grant from the then Director (Prof. Animesh Biswas) of the NIT-Rourkela to SKP.

## Conflict of Interest

None

## Notes

### Competing Interest Statement

The authors have declared no competing interest.

