## Supplementary material for "Modulation of H3K4 trimethylation by KDM5A and MLLs impacts metabolic adaptability in prostate and cervical cancer cells": Yes

### Supplementary Information

**siRNAs used in this study:**
KDM5A siRNA: Thermo – siRNA ID – s11836 (Ambion silencer select siRNA)
Sequence: Sense – GCGAGUUUGUUGUGACAUUtt; Antisense – AAUGUCACAACAAACUCGCca
MLL1 siRNA: Sigma – siRNA ID – SASI_Hs01_00090459
Negative control: Thermo – catalog no – 4390843 (Silencer select negative control siRNA)

**Drugs used:**
Geraniol: Sigma 163333, Embelin: SC201555, Gallic acid: Fluka 91215, Stigmasterol: SC281156 and curcumin: Sigma C1386. All these compounds were dissolved in DMSO.

**RT-PCR primers used in this study:**

**1. Hexokinase 2**

FP- AAGTGCAGAAGGTTGACCAG

RP- CCAAGCCCTTTCTCCATCTC

**2. Phosphofructokinase (PFKFB3)**

FP-AAAAGCCTCGCATCAACAGC

RP-TCCGGGAGCCTTTCATGTTT

**3. Pyruvate Kinase M2**

FP- ATGCGGAGACCATCAAGAAT

RP- GTTCGGATCTCAGGTCCTTT

**4. Pyruvate Dehydrogenase**

FP- CTGAAGAGGCGCTTTCACTG

RP- TCTCATCTCTTTCCAGCTCCTC

**5. Alpha Ketoglutarate Dehydrogenase (E1)**

FP- GCACAGTCCCTGGTAGAAGC

RP- ACATGGTGCCCTCGTATCTG

**6. NDUFS3**

FP- GTGTTCAAGGCAGCCAACTG

RP- AAAGGATGTCCCTCGAAGCC

**7. NDUFV2**

FP- CGAGTTCTTCAGCGGACTCA

RP- GTCCACTGGGAAGGATCAGG

**8. SDHD**

FP- GCTTCGAACTCCAGTGGTCA

RP- AATGGTGGCTCGGTGACAAG

**9. CYC1**

FP- GCTTCGCGGGGTAGTGTTG

RP- CACTGCCTGAGGTGTCCGTA

**10. COX4|1**

FP- CGCTCGTTATCATGTGGCAG

RP- GGGGTTCACCTTCATGTCCA

**11. ATP5F1A**

FP- CGAATTTCAGTGCGGGAACC

RP- GGTTTTCCCAGTCTGTCGGT

**12. ATP5F1B**

FP- GGTCCTGAGACTTTGGGCAG

RP- TGGAGCCTCAGCATGAATGG

**13. DRP1**

FP- ACTGGAATTGTCACCCGGAG

RP- TCTGCTTCCACCCCATTTTCT

**14. MFN1**

FP- AGCGGCTTTCCAAGCCTAAT

RP- CTTTCCATGTGCTGTCTGCG

**15. MFN2**

FP- TCTCCCGGCCAAACATCTTC

RP- ACCAGGAAGCTGGTACAACG

16. **UCP2**

FP- AGCCCACGGATGTGGTAAAG

RP- CTCTCGGGCAATGGTCTTGT

17. **UCP3**

FP- CCCTCCTTTTTGCGTTTGGG

RP- ACGGTGATTCCCGTAACATCT

18. **SOD1**

FP- TGGCCGATGTGTCTATTGAA

RP- CTTCATTTCCACCTTTGCCC

19. **Catalase**

FP- ACCAAGGTTTGGCCTCACAA

RP- TGCTTGGGTCGAAGGCTATC

20. **GPX1**

FP- AACCAGTTTGGGCATCAGGA

RP- AGAGCATGAAGTTGGGCTCG

21. **Glutathione reductase**

FP- TGAAGTTCTCCCAGGTCAAG

RP- GGCAGTCAACATCTGGAATC

22. **Glucose 6-P Dehydrogenase**

FP- TAGGCTGGAACCGCATCATC

RP- TGCGGTAGATCTGGTCCTCA

23. **Lactate Dehydrogenase A**

FP- CCAACATGGCAGCCTTTTCC

RP- TTCTCCCTCTTGCTGACGTG

**Antibodies used in this study:**

The antibodies against KDM5A and MLL2 was purchased from Abcam; PFK1, IDH3A, ATP1F1, SOD1, Catalase, G6PD, MFN2 and DRP1 were procured from Santacruz and H3K4me3 was procured from Invitrogen (catalogue numbers are listed below).

KDM5A (ab70892)
KMT2b (ab56770)
Anti-PFK-1 Antibody (G-11): sc-166722.
Anti-IDH3A Antibody (A-10): sc-398021.
Anti-ATPIF1 Antibody (A-3): sc-271614.
Anti-Superoxide Dismutase 1/SOD1 Antibody (G-11): sc-17767.
Anti-catalase Antibody (H-9): sc-271803.
Anti-G6PD Antibody (G-12): sc-373886.
Anti-DRP1 Antibody (6Z-82): sc-101270.
Anti-Mfn2/Mitofusin 2 Antibody (F-5): sc-515647.
H3K4me3 (Invitrogen MA5-11199)

**Chip buffer composition and primers used:**

ChIP lysis buffer (1% SDS, 10 mM EDTA, 50 mM Tris-HCl, pH 8, and protease inhibitors)
ChIP dilution buffer - (0.01% SDS, 1% Triton X-100, 1.2 mM EDTA, 16.7 mM Tris-HCl, pH 8, 167 mM NaCl, and protease inhibitors)
low-salt buffer- (0.1% SDS, 1% Triton X-100, 2 mM EDTA, 20 mM Tris-HCl, pH 8, 150 mM NaCl)
high-salt buffer -(0.1% SDS, 1% Triton X-100, 2 mM EDTA, 20 mM Tris-HCl, pH 8, 500 mM NaCl)
LiCl buffer (250 mM LiCl, 1% NP-40, 1% deoxycholate, 1 mM EDTA, 10 mM Tris-HCl, pH 8)
TE buffer (10 mM Tris-HCl, pH 7.6, 1 mM EDTA)
Elution buffer - (0.1 M NaHCO3, 1% SDS) (freshly made)

% INPUT was calculated as follows:

% INPUT = 100* 2x-y

(Where X is the ct value for Input adjusted for dilution factor and Y is the ct value for the IP samples.

Catalase ChIP primer:

Catalase (137 bp)

F-CCCGAAGGTCCGTTTAGAA

R- GCGAATGTAAAAGTCCGTCT

#### Supplementary data

##### Supplementary figure legends:

**Supplementary figure S1:** MLL1 expression profile in CESC and PRAD tumors was less compared to normal tissues as observed from Gepia2 database gene expression profiling data with bars representing median expression of gene in transcripts per million.

**Supplementary figure S2:** MLL1 knockdown using siRNA was confirmed by immunoblotting using MLL1 antibody, followed by staining with Alexa647. Images shown are one of the 3 independent repeats. Student's 't' test was used to test significance; *$P \leq 0.05$, **$P \leq 0.01$, ***$P \leq 0.001$ and ns is non-significant i.e., $P > 0.05$.

**Supplementary figure S3**: Box plots indicating the expression of G6PD in cervical (CESC) and prostate (PRAD) tumors compared to their respective normal tissues was obtained using UALCAN database. **(a).** G6PD was observed to be highly enhanced in cervical cancer patients (n=305) compared to normal (n=3), but unaltered in prostate cancer patients **(b).**

**Supplementary figure S4:** MLL1, MLL2, H3K4me3 and KDM5A ChIP-seq data. Only KDM5A, MLL2 and H3K4me3 shows an overlap of occupancy on PFKM (A), OGDH (B), NDUFA4 (C), COX4I1 (D), RPIA (E) and MFN2 (F) promoters, but MLL1 did not show any enrichment on these promoters.

**Supplementary figure S5**: The graphs show the cellular growth changes in HeLa and HaCaT cell lines following administration of geraniol, embelin, gallic acid, stigmasterol and curcumin; as assessed by MTT assay.

##### Supplementary table legends:

**Supplementary table T1:** The data presents correlation coefficient with respective p- and q values obtained from cBioPortal to understand correlation between different metabolic signature genes with KDM5A, as obtained from TCGA cervical cancer patient datasets.

**Supplementary table T2:** A negative correlation between 56 metabolic signature genes with KDM5A was observed in prostate cancer patient samples obtained from TCGA, the table depicts correlation coefficient with respective p- and q values obtained from cBioPortal.

**Supplementary table T3:** Individual mRNA expression values of genes plotted in figure 2.a, as retrieved from cBioPortal. These values correspond to the 294 CSCC patient samples obtained from TCGA.

**Supplementary table T4:** Statistical information pertaining to Z-scores and correlation coefficient of genes plotted in figure 2.a

**Supplementary table T5:** Individual mRNA expression values of genes plotted in figure 2.b, as retrieved from cBioPortal. These values correspond to the 493 PRAD patient samples obtained from TCGA.

**Supplementary table T6:** Statistical information pertaining to Z-scores and correlation coefficient of genes plotted in figure 2.b

**Supplementary table T7**: Correlation between different metabolic signature genes with MLL1 obtained from TCGA cervical cancer patient dataset. The table presents data on correlation coefficient with respective p- and q values obtained from cBioPortal.

**Supplementary table T8:** Correlation between different metabolic signature genes with MLL1 obtained from TCGA prostate cancer patient dataset with spearman's coefficient, p- and q-values.

**Supplementary figures:**

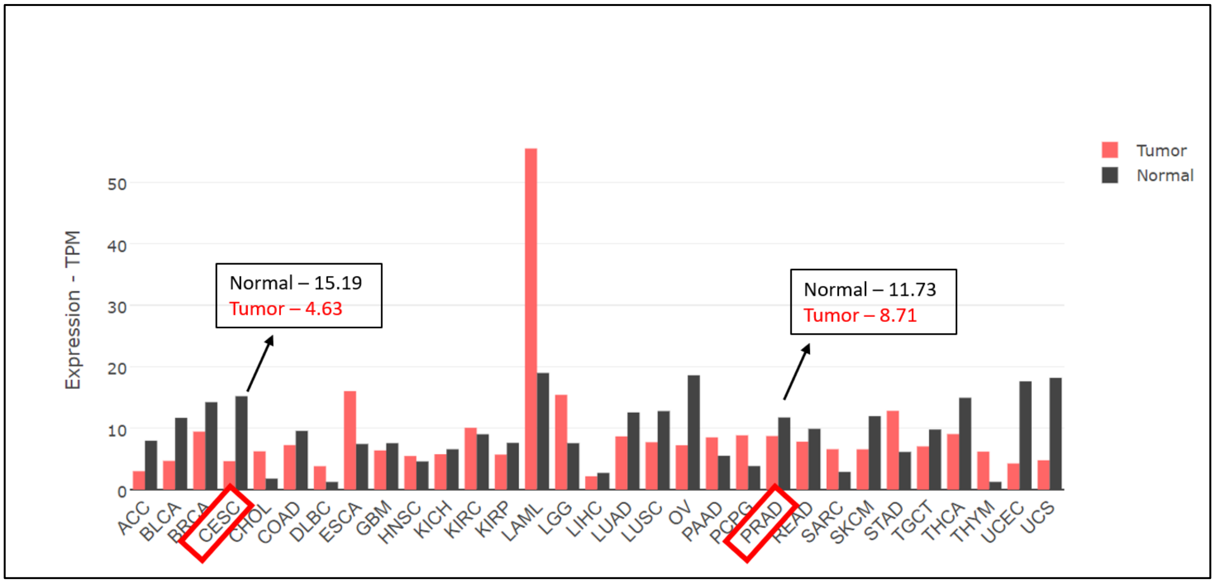


**Supplementary figure S1**

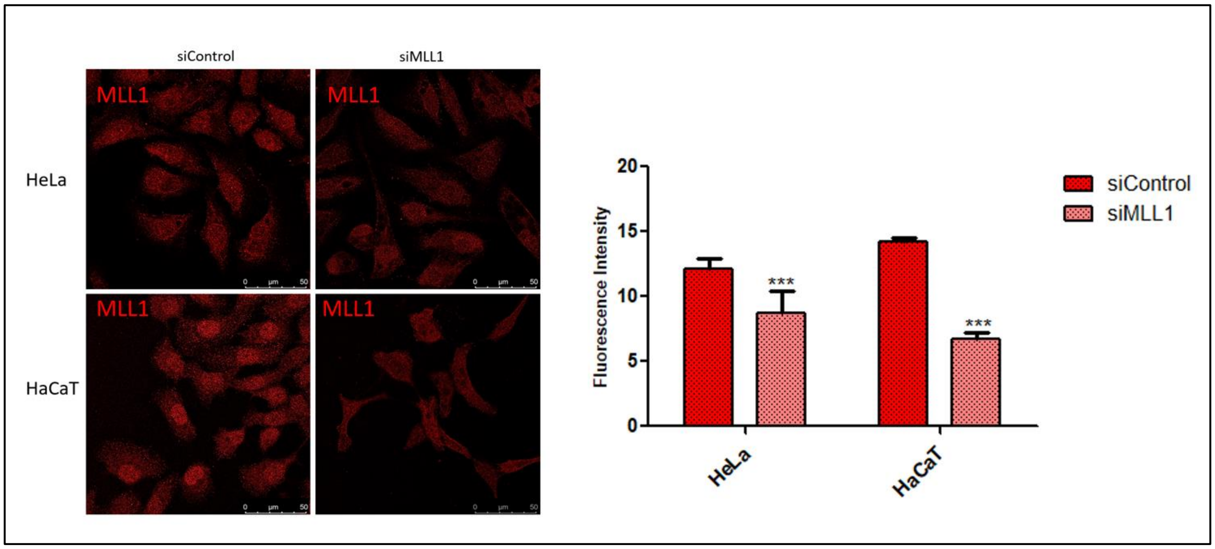


**Supplementary figure S2**

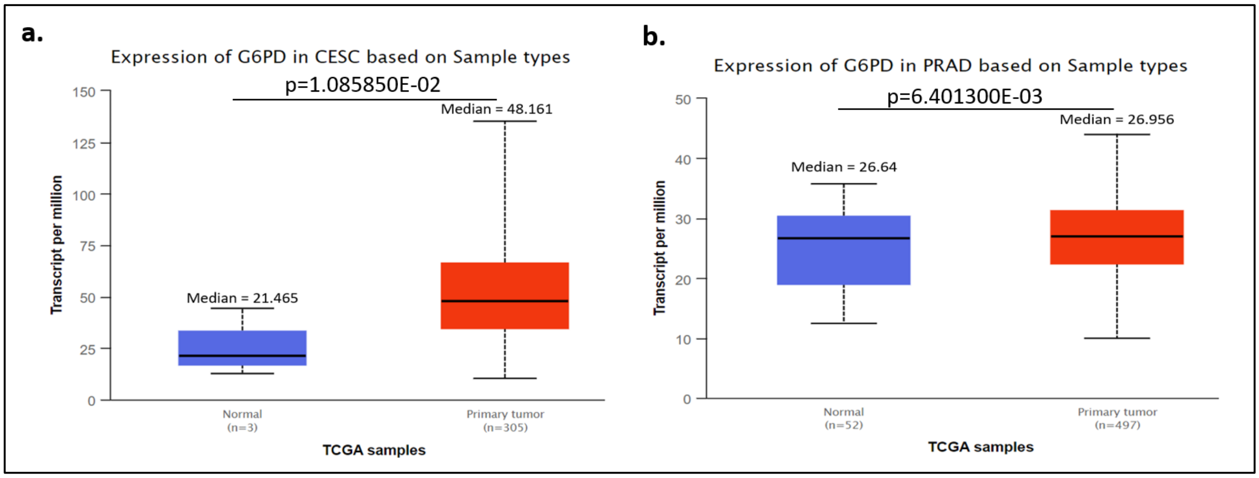


**Supplementary figure S3**

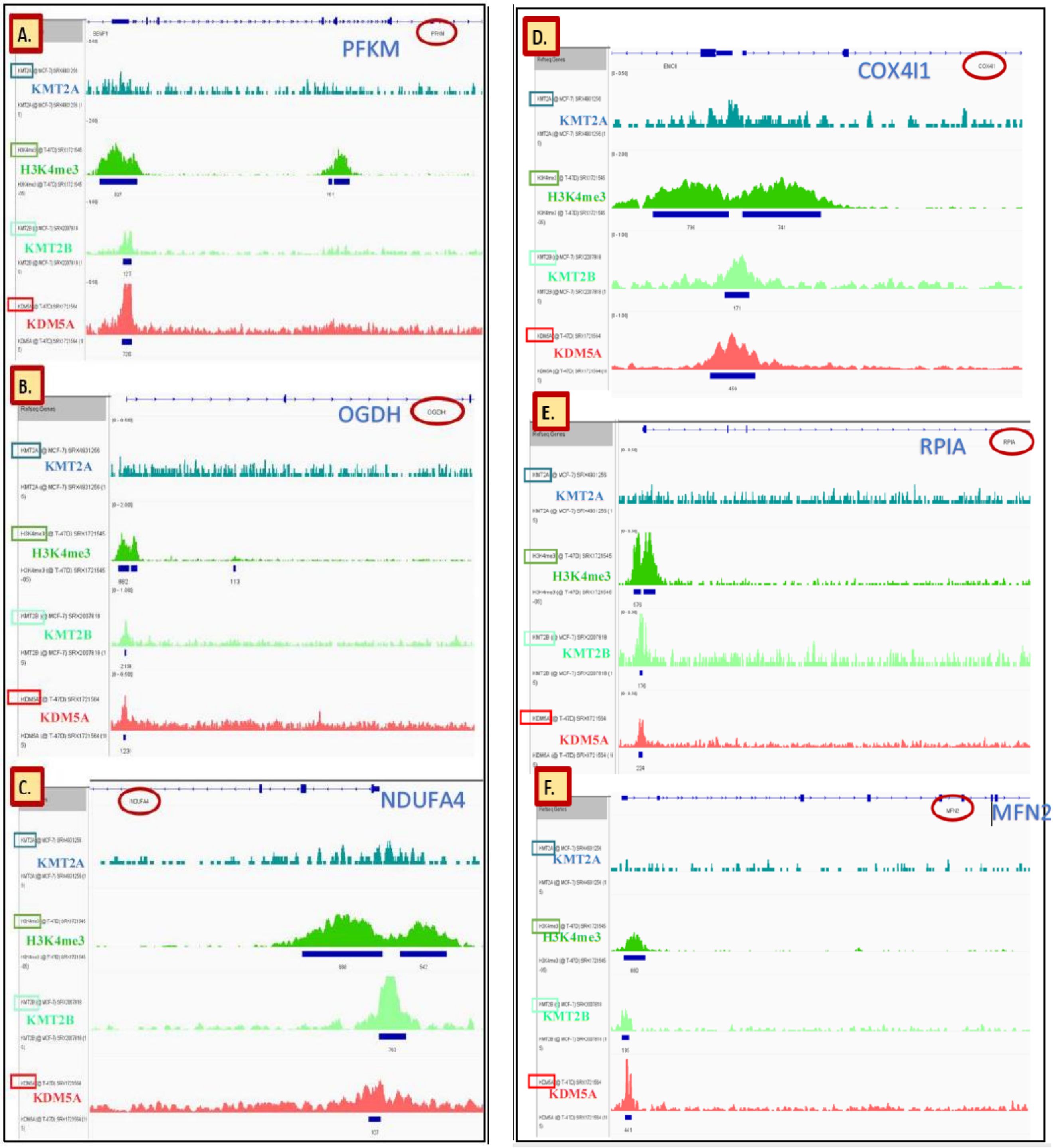


**Supplementary figure S4**

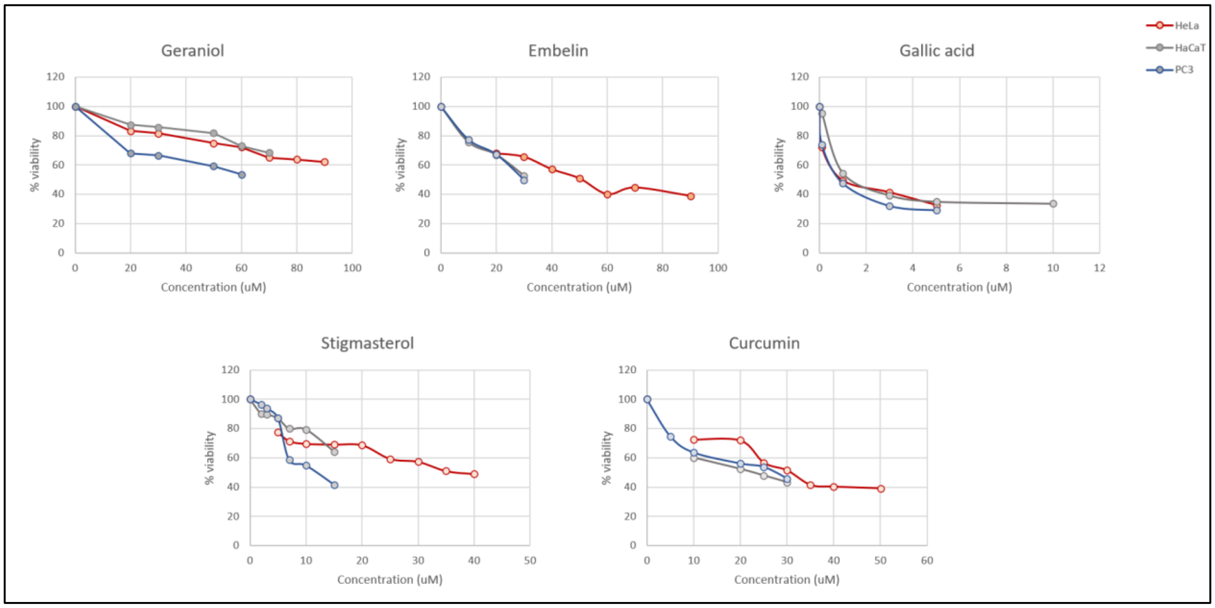


**Supplementary figure S5**
